# Multiple polyploidizations in *Arabidopsis lyrata* stabilized by long-range adaptive introgression across Eurasia

**DOI:** 10.1101/2024.08.27.609292

**Authors:** Alison D. Scott, Uliana Kolesnikova, Anna Glushkevich, Laura Steinmann, Nikita Tikhomirov, Ursula Pfordt, Magdalena Bohutínská, Robin Burns, Alexey P. Seregin, Filip Kolar, Roswitha Schmickl, Polina Yu. Novikova

## Abstract

Abundance of polyploidy varies across lineages, evolutionary time and geography, suggesting both genetics and environment play a role in polyploid persistence. *Arabidopsis lyrata* appears to be the most polyploidy-rich species-complex in the *Arabidopsis* genus, with multiple origins of autotetraploidy. This is revealed by genomic data from over 400 samples across Eurasia. We found over 30 previously undescribed autotetraploid populations in Siberia with a minimum of two separate origins, independent of those previously reported in Central Europe. The establishment of Siberian tetraploids is mediated by meiotic adaptation at the same genes as in European tetraploid *A. lyrata* and *Arabidopsis arenosa,* despite high divergence and geographical separation. Haplotype analysis based on synthetic long-read assemblies supports the long-range introgression of adaptive alleles from the tetraploid interspecific pool of European *A. lyrata* and *A. arenosa* to tetraploid Siberian *A. lyrata*. Once evolved, adaptation to polyploidy promotes the establishment of new polyploid lineages through adaptive inter– and intraspecific introgression.

## Introduction

Polyploidization cycles are recurrent and abundant with non-random distribution in space, through time, and across the Tree of Life. Polyploid frequency has a latitudinal gradient: more polyploids are observed in colder regions closer to the poles^1,2^, which is paralleled with dating of ancient WGD events at environmentally harsh times^3^. Whole genome duplications are recurrent; both monocot and dicot lineages experienced multiple polyploidy events detected in the period between 75 and 55 Mya^4^, a time range of drastic climatic changes. Polyploidy is more abundant in plants^3,5^ compared to animals^6^, in Angiosperms vs Gymnosperms^7^, and in Amphibia vs Mammals^6,8^. What explains the abundance of polyploids in certain lineages, their emergence at certain times and climates? The explanation of the non-random distribution of polyploidy must lie in the balance between the birth and death rate of whole genome duplications, which differs depending on the context. For example, the environment, a function of time and space, can both trigger polyploidy through unreduced gamete production^9–11^ and impact extinction rates^3^. Genetic background also affects polyploid birth rates directly through production rates of unreduced gametes^12–14^, and can provide a route to frequent polyploid emergence.

The emergence of polyploids does not guarantee their survival. Neopolyploid establishment includes adaptation to external environments^15,16^, but adaptation to the new internal environment is even more critical. Whole genome duplication in an individual (or single lineage; autopolyploidy) results in multiple sets of homologous chromosomes. Challenges in executing faithful segregation of the doubled chromosomes can hinder the survival of neopolyploids^17,18^. Artificially induced polyploids have reduced fertility and show high levels of aneuploidy in subsequent generations^19–22^, suggesting that natural polyploids have evolved adaptations to the polyploid state^23,24^ or may be established only from certain preadaptive combinations of alleles in the diploid pool^22^.

The genus *Arabidopsis* is a prime example of the non-random nature of polyploidy. Multiple lineages are either fully polyploid (*A. suecica*, *A. kamchatica*) or harbor polyploid populations (*A. arenosa*, *A. lyrata*). *Arabidopsis* polyploids emerged in times of climatic upheaval^25^. *A. arenosa* polyploids have been described to have a single geographical origin in the western Carpathian Mountains and spread across Europe after a whole-genome duplication^26,27^. Establishing tetraploid *A. arenosa* involved polygenic adaptation of reproductive machinery, including pollen tube growth^28^ and meiosis. The former is probably to restore the ion homeostasis, the latter is to prevent entanglement between three or more chromosomal copies and ensure faithful segregation^18,20,23^.

We focus our study on the most Northern *Arabidopsis* species with a wide geographical distribution – *Arabidopsis lyrata*. Multiple tetraploid *A. lyrata* populations have been described in Europe in several areas in the Czech Republic and Austrian Alpine foothills^29–32^. Tetraploid *A. lyrata* populations in Central Europe form three distinct lineages occupying three distinct regions^33^, which may suggest multiple origins of polyploidy in contrast to *A. arenosa*. Diploid and tetraploid *A. lyrata* and *A. arenosa* have overlapping, yet non-sympatric ranges in Europe, and while diploids are reproductively isolated^34^ and do not show signs of ongoing interspecific gene flow^29,32,35,36^, tetraploid *A. lyrata* and *A. arenosa* often hybridize^30,32,37–40^. Interestingly, interspecific gene flow between European tetraploid *A. lyrata* and *A. arenosa* has been adaptive: several introgressed alleles of meiotic genes from *A. arenosa* are also under selection in tetraploid *A. lyrata*, as they increase the fitness of tetraploids by stabilizing chromosomal segregation^39,41^.

There are indications of a polyploid population in Yakutia^25,42^, suggesting that ploidy variation could still be massively underestimated in this species due to a lack of extensive sampling, particularly in Northern Eurasia. Despite the wide distribution of *A. lyrata* across the Northern Hemisphere^43–46^, a large part of the described range from Eastern Europe to Eastern Siberia lacks genetic data. We generate a whole-genome re-sequencing dataset from this understudied region and ask if polyploids are abundant there. Here we report multiple new tetraploid *A. lyrata* populations in the areas of White Sea shore, Polar Ural, and Central Siberia. We describe the nature, origins and relationships of these WGDs. We ask what explains the abundance of polyploids in *A. lyrata* in these remote geographical regions since the distribution of previously described adapted polyploids is limited to European *A. arenosa* and *A. lyrata*. We investigate if any of the alleles known to stabilize polyploid meiosis could reach from Europe to Eastern Siberia or adaptation to polyploidy is purely convergent.

## Results

### Tetraploid *A. lyrata* is abundant in Siberia

We assembled a dataset of whole-genome sequences from several sources. First, we sequenced 116 herbarium samples from the Moscow University Herbarium^47^, spanning a wide geographical area from the White Sea to the Bering Strait. Second, we sequenced several individuals per population (283 samples from 32 locales) grown from seeds collected during expeditions to the White Sea, Polar Urals, Gydan Peninsula, Putorana Plateau, Lower Lena and Amur (Figure 1a, Supplementary Figure 1, Supplementary Data 1). Third, we included whole-genome sequencing data of samples from different *A. lyrata* lineages which have been previously published^39,45,48–51^. All re-sequencing data were mapped to the reference genome of the self-compatible Siberian *A. lyrata* NT1 strain^52^.

**Figure 1.**
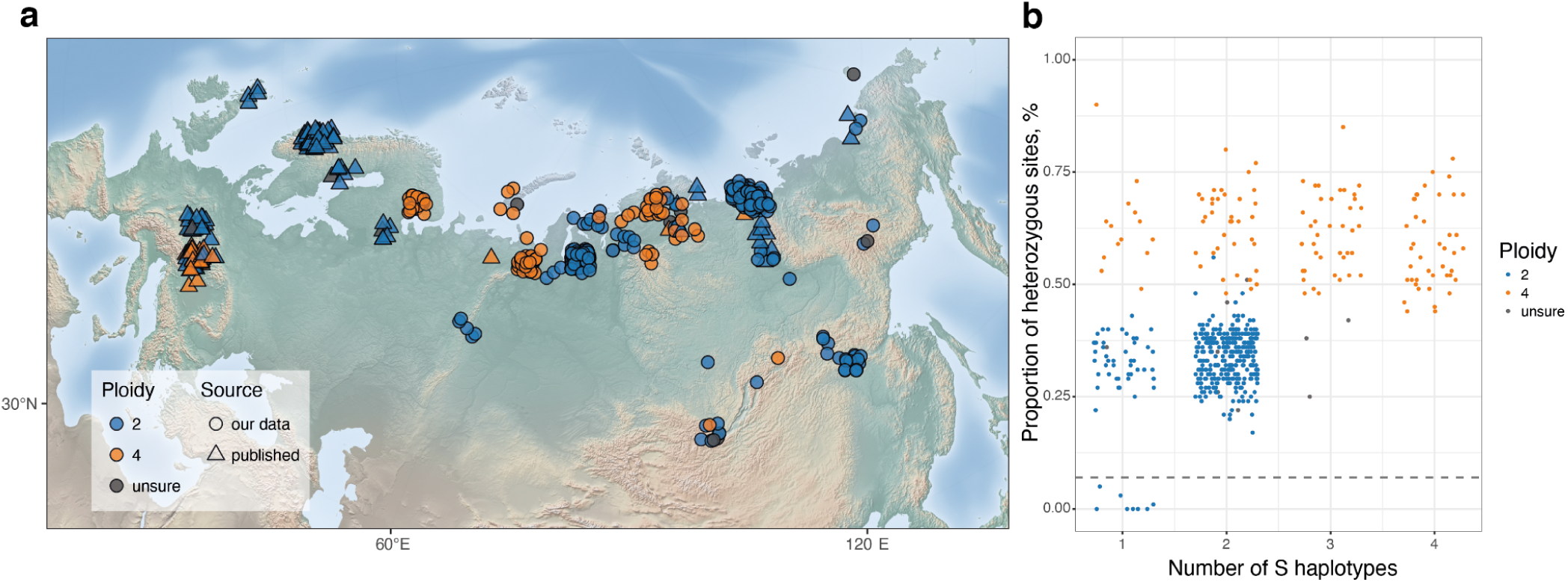
Distribution of diploid and tetraploid *A. lyrata* across Eurasia. (**a**) Geographical distribution of *A. lyrata* samples used in this study. For better visualization in cases of overlaps we randomly shifted the points within 100 km from the actual sampling location, all true coordinates can be found in the Supplementary Data 1. Newly found tetraploid *A. lyrata* populations are represented in orange circles. (**b**) Relationships between proportions of heterozygous biallelic SNPs and the number of different *S*-alleles in each individual shown in (a). Tetraploid individuals (orange dots) show a higher proportion of heterozygous sites and up to four different *S*-alleles. Diploid individuals (blue dots) show a lower proportion of heterozygous sites and up to two different S-alleles; individuals with one *S*-allele below the dashed gray line correspond to recently described Siberian selfing *A. lyrata* lineage with nonfunctional *S*-locus^62^. Herbarium individuals with ambiguous assignments of ploidy between two inference methods are shown in gray dots.

We used a statistical framework based on relative allelic counts implemented in nQuire software^53^ to get a first inference of ploidy from sequencing data (See *Methods*). We initially called variants for all samples as diploids. The proportion of heterozygous sites in tetraploid individuals was generally higher than in the diploids (Figure 1b, Supplementary Figure 2). To complement the initial inference, we genotyped and counted *S*-alleles in each sample using NGSgenotyp^54^. Due to the extreme divergence and diversity of *S*-alleles controlling self-incompatibility in *Arabidopsis*^55–61^, it was often possible to find up to two different *S*-alleles in diploid and up to four different *S*-alleles in tetraploid individuals. These independent assessments of ploidy levels allowed us to infer ploidy for most of the samples (Figure 1b). Finally, we directly counted chromosome numbers and measured genome sizes in the live samples grown in the greenhouse from seeds collected in the field (Supplementary Figure 1, Supplementary Data 2) and confirmed our bioinformatic inferences. When bioinformatic inference yielded unclear results and cytogenetics was not possible (i.e. for herbarium specimens), we indicate the ploidy as “unsure” and exclude these accessions from downstream analyses (Figure 1b, Supplementary Data 1). Also, while tetraploid and diploid populations sometimes occur nearby, we found no evidence of mixed-ploidy populations: all individuals sampled from the same population had consistent ploidy levels (Supplementary Data 1). Following the assignment of ploidy by heterozygosity, *S*-allele genotype, and cytological confirmation, we re-called variants according to the inferred ploidy state using the designated ploidy-aware approach in GATK.

Our live collections included confirmed self-compatible and self-incompatible diploid *A. lyrata* individuals, which differ in the levels of heterozygosity (Kolesnikova et al. 2023) (Figure 1b, Supplementary Figure 2). Tetraploid individuals had a significantly higher proportion of heterozygous sites compared to diploid selfers and outcrossers, and up to four different *S*-alleles in each individual, which points to their outcrossing nature (Figure 1b, Supplementary Figure 2), confirmed by greenhouse observations. Nucleotide diversity (π) calculated on putatively neutral four-fold degenerate sites is comparable between diploid and tetraploid (∼0.005) *A. lyrata* geographic populations (individuals sampled in the same collection site) (Supplementary Table 1), showing no evidence of genetic bottleneck in tetraploids.

### Independent autopolyploidization events of A. lyrata in Northern Ural and Central Siberia around LGM

To characterize the population structure of *A. lyrata* across Eurasia, we calculated pairwise genetic distances from the four-fold degenerate sites as a proxy for nearly neutral sites and built a network (Figure 2a), showing that the newly described tetraploids cluster within two distinct groups, separate from the Central Europe tetraploids. Northern Ural (NU, highlighted in pink, Figure 2a) populations sampled around the White Sea and in Polar Ural mountains form a cluster containing diploid and tetraploid individuals. This NU cluster is genetically close to the West Siberian cluster (WS, highlighted in blue, Figure 2a) comprised purely of diploid populations sampled from Taz Estuary. Central Siberian cluster (CS, highlighted in yellow, Figure 2a) also contains both diploids and tetraploids and, in turn, is closer to diploids from the lower Lena river area in East Siberia (ES, highlighted in orange, Figure 2a). Tetraploids are genetically closer to their spatially close diploid populations from their corresponding clusters (NU 4x to NU 2x and CS 4x to CS 2x, CE4x) than each other, suggesting their independent origin.

**Figure 2.**
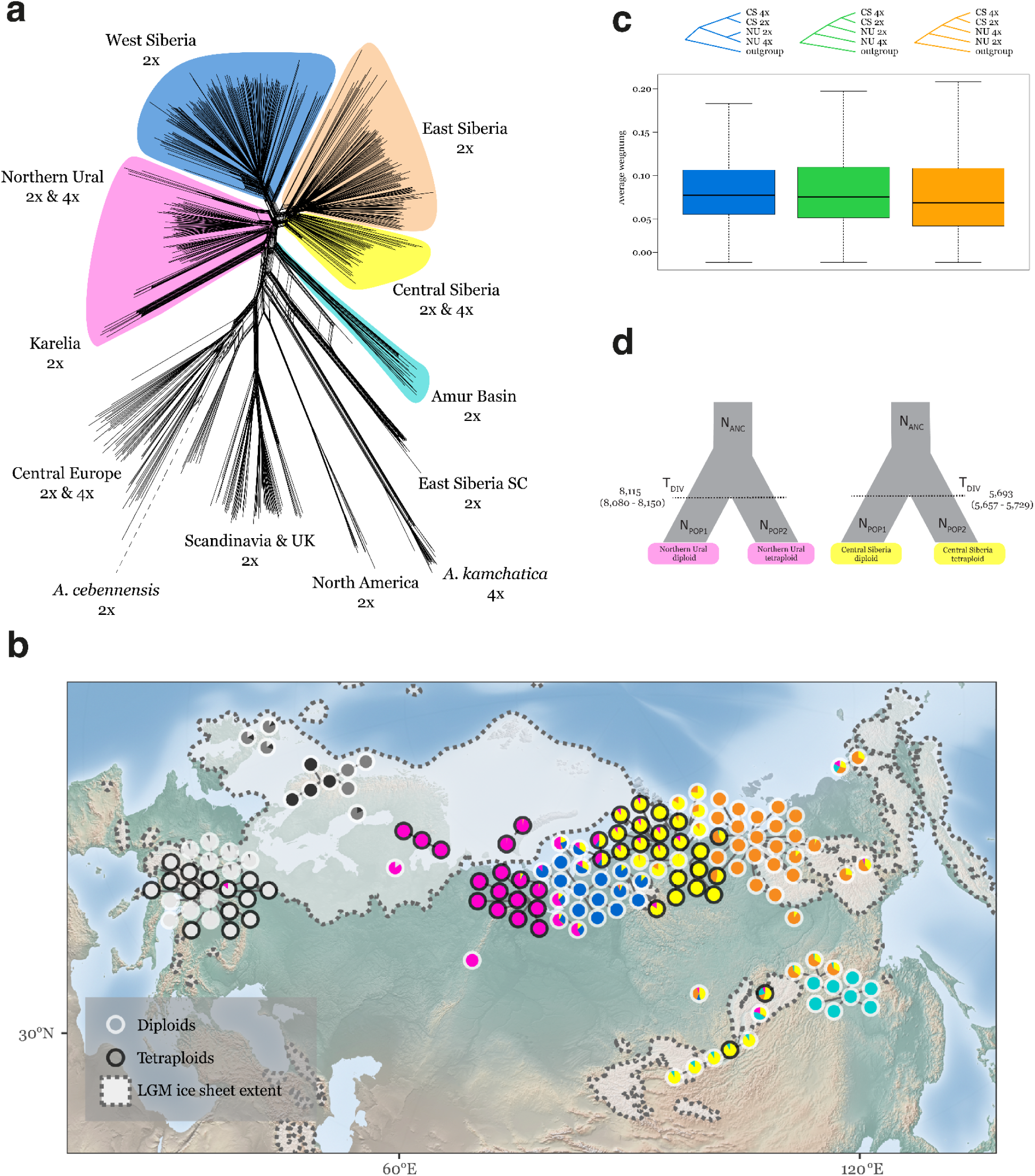
Population structure of *A. lyrata* diploid-tetraploid species complex. (**a**) Network representation of genetic pairwise distances between individuals. *A. lyrata* accessions form several clusters outside of Central Europe, named here as Northern Ural (NU; pink, diploid/tetraploid), West Siberian (WS; blue, diploid), Central Siberian (CS; yellow, diploid/tetraploid), East Siberian (ES; orange, diploid), and Amur Basin (AB; turquoise, diploid). Note that while network clusters may contain multiple ploidy levels, geographic populations are not mixed ploidy. Newly described autotetraploids are found in two distinct clusters (NU/pink and CS/yellow), distinct from Central European autotetraploids (gray) and allotetraploid *A. kamchatica*. (**b**) Admixture across the Eurasian *A. lyrata* distribution. Colors correspond to labeling in (**a**) and reflect average admixture proportions per sampling site. Tetraploid accessions are found across the range (denoted by black outlines). Extent of the ice sheet during the Last Glacial Maximum^69^ shown in white. (**c**) The three highest weighted topologies of populations show that diploid and tetraploid Northern Ural populations form a clade; the same for diploid and tetraploid Central Siberian populations. (**d**) The estimates of the time of origins for the two tetraploid lineages as divergence time estimates from the closest diploid progenitors using Fastsimcoal2^70^.

Using population-level data, we estimated site frequency spectra for the two newly discovered tetraploid *A. lyrata* lineages (NU 4x and CS 4x) separately and compared them to theoretical expectations for auto– and allo-polyploids. In both cases, the observed site frequency spectra of Siberian *A. lyrata* tetraploids are consistent with autopolyploidy, indicated by an absence of sites at an intermediate allele frequency (Supplementary Figure 3). Allotetraploid populations exhibit a pronounced peak at the intermediate frequency when mapped to a diploid reference due to subgenome divergence, while autotetraploid populations do not^63^. Allotetraploid *Arabidopsis kamchatica*, originating through hybridizations of Siberian *A. lyrata* and *Arabidopsis halleri*^62,64,65^, is also included in the network and clusters separately from NU and CS autotetraploid *A. lyrata* (Figure 2a).

We then used TreeMix^66^ to describe relationships among diploid populations. The maximum likelihood model (Supplementary Figure 4a) groups Karelia, Northern Ural, Western Siberia, Central Siberia, Eastern Siberia, and Amur Basin together with suggested migration edge between Karelia and Central Europe, while UK and Scandinavian populations grouped with Central European populations. In the maximum likelihood model including all diploid and tetraploid populations, neither European nor Siberian tetraploids grouped together: in both cases instead grouping with geographically close diploids. (Supplementary Figure 4b). Clustering of individuals using ADMIXTURE^67^ with the optimal number of groups K=8 (Supplementary Figure 5) assigned tetraploid and diploid NU populations into a single cluster (pink, Figure 2b with high levels of admixture with diploid WS populations (blue, Figure 2b); tetraploids and diploids from CS were also assigned into a single cluster (yellow, Figure 2b) with high levels of admixture with diploid ES cluster (orange, Figure 2b). We used topology weighting with Twisst^68^ to further explore the relationships between *A. lyrata* lineages when variation in topologies across the genome is taken into account (Figure 2c, Supplementary Figure 6). The top three highest-weighted topologies are consistent with the independent origin of the two autotetraploid lineages. Northern Ural tetraploids and diploids form a monophyletic clade in the most frequent topology, as do Central Siberian tetraploids and diploids.

To estimate the timing of tetraploid origin in Northern Ural and Central Siberia, we used a demographic modeling approach based on fitting simulated and observed site frequency spectra^70^. For each pair of diploid-tetraploid populations (NU 2x and NU 4x, CS 2x and CS 4x) we tested a simple divergence model and a simple model with unidirectional gene flow from diploids into tetraploids, assuming gene flow can be realized through unreduced gametes formed by diploids. Likelihood comparisons and AIC tests suggested that the gene flow model better fit the data in both cases. For the origin of Northern Ural tetraploids, we estimated 8,115 generations ago (95% CI 8,080-8,150), which we date at roughly 16 Kya (assuming 2 year generation time), and for the Central Siberian tetraploids we estimated an origin at 5,693 generations ago or approximately 11 Kya (95% CI 5,693-5,729) (Figure 2d, Supplementary Table 2).

### Tetraploid-specific adaptive introgression of genes involved in meiotic stabilization

While topologies consistent with the population tree and the network analysis were the most frequent (Figure 2c), a smaller but considerable fraction indicated introgression between the Northern Ural and Central Siberian tetraploids (topologies 4,5,6, Supplementary Figure 6). We combine the topology weighting results for all introgression topologies (4,5,6) and show them along the chromosomes (Figure 3a, top panels). In total, 11 windows with a high weighting of gene flow topologies are apparent. Comparing these window boundaries with annotation, we identified 37 gene models (Supplementary Data 4) located within introgression windows. These genes are enriched for gene ontology terms related to meiosis and chromosome organization (Figure 3b, Supplementary Data 4). Performing the same analysis now including Central European and Northern Ural diploid and tetraploid populations, we find striking overlap of the introgression windows found in Northern Ural and Central Siberian comparison (Figure 3a, bottom panels). For each gene in introgression windows, we look at SNP frequencies across diploid and tetraploid lineages including Central Europe, Northern Ural and Central Siberia (Supplemental Figure 7). We find SNP sharing across the entire geographical range: in some cases SNPs appear to be tetraploid-specific (e.g. *ASY3*, *MAU2*, *PDS5*, and *ZYP1b*) and other SNPs are shared among diploids and tetraploids (e.g in *CYCA2;3* and *PAPP2C*).

**Figure 3.**
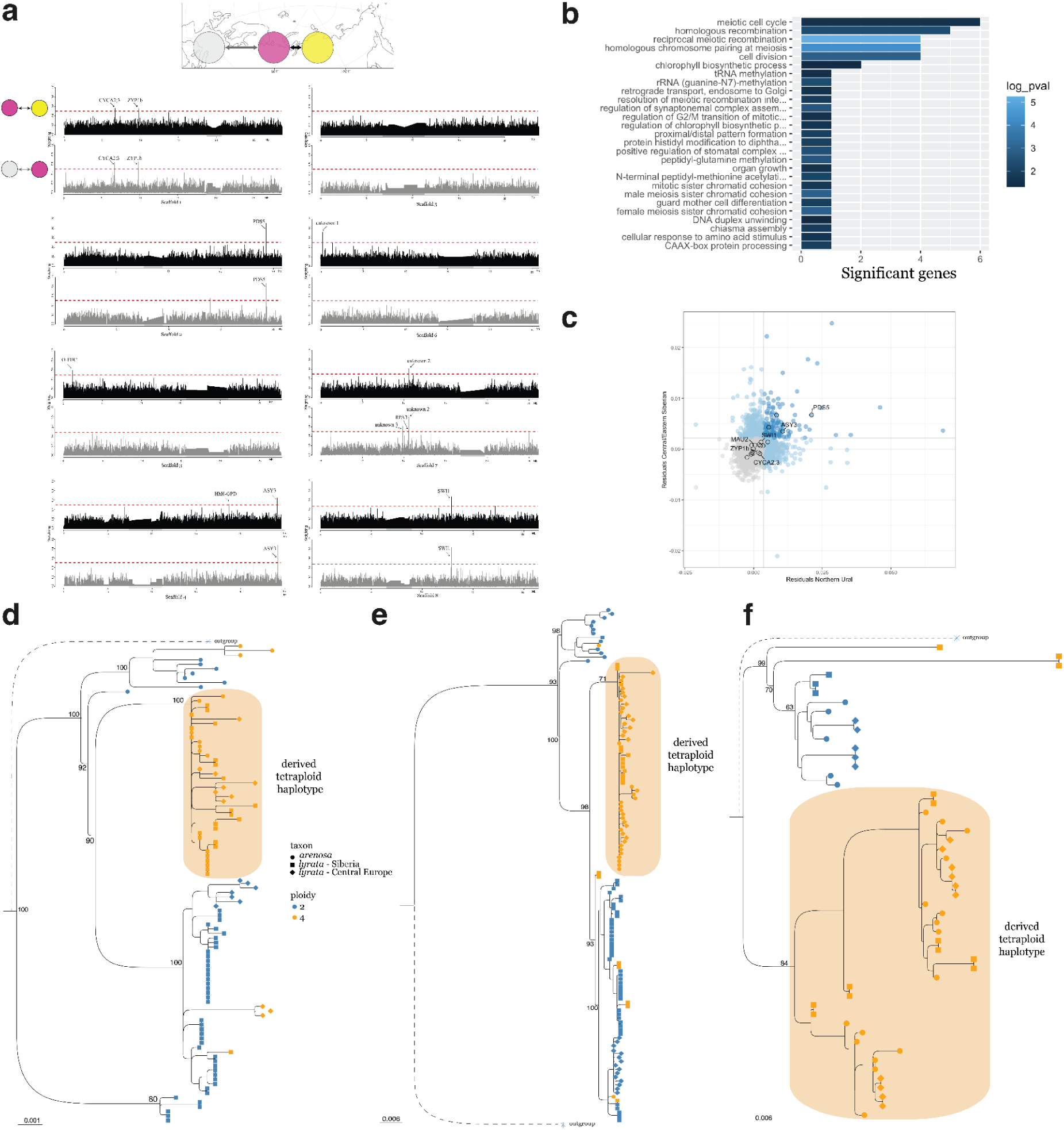
Genes present in introgression topologies. (**a**) Distribution of introgression topology windows and their weighting across the genome, per scaffold. Height of peaks indicates weight, and introgression blocks are annotated. (**b**) Results of GO term enrichment for genes within introgression windows, with number of significant genes indicated on the x axis and log p-value by color of bars. (**c**) Scatter plot of residuals from linear model fit to nucleotide diversity (π) within tetraploids and divergence (dxy) between diploids and tetraploids. Nucleotide diversity is calculated per-gene, so that each dot represents a single gene. Outlier genes colored light blue for each comparison (Northern Ural diploids and tetraploids on x axis; Central Siberian diploids and tetraploids on y axis), darker blue indicates outliers in both comparisons. ASY3 and PDS5 annotated within outliers. (**d, e, f**) gene trees for haplotypes from *A. lyrata* in Siberia, *A. lyrata* in Central Europe, and *A. arenosa*. Each tip is a haplotype, orange tips are haplotypes from tetraploid individuals, and blue tips from diploids. Shape indicates taxon: squares for *A. lyrata* in Siberia, diamonds for *A. lyrata* in Central Europe, and circles for *A. arenosa*. (**d**) Haplotype tree for *PDS5* (**e**) Haplotype tree for *ASY3* (**f**) Haplotype tree for *ZYP1b*.

The strong enrichment of specific gene ontology terms (Figure 3b) suggests introgressed alleles are undergoing positive selection. To verify this, we calculated nucleotide diversity (π) per gene within Northern Ural tetraploids and between Northern Ural diploids and tetraploids and between Central Siberian diploids and tetraploids using piawka (https://github.com/novikovalab/piawka). Under neutrality, these values should be correlated, and outliers should indicate selection (Supplementary Figure 8). Four introgressed genes deviated from neutral expectations and instead fall among outliers in both comparisons (Figure 3c), concluding that genes prone to introgression between the Northern Ural and Central Siberian tetraploids are also under selection in both tetraploid lineages.

To validate haplotype sharing across the range, we obtained full haplotype sequences from synthetic long reads (single-tube long fragment reads; stLFR), PacBio long-read assemblies (see Methods), and published haplotype sequences of Sanger sequenced PCR products^23,71^. We focused on three introgressed genes (*ASY3*, *PDS5*, and *ZYP1b*) with substantial published sequence data across *Arabidopsis* genus allowing haplotype-scale alignments for further analyses. We aligned assembled haplotypes from diploid and tetraploid *A. arenosa* and *A. lyrata* and estimated phylogenetic trees. Tetraploid-specific haplotypes from *A. arenosa*, Central European, Northern Ural and Central Siberian *A. lyrata* form a monophyletic clade at these loci (Figure 3d-f). This corresponds well with SNP frequency heatmaps at the same loci (Supplemental Figure 7).

## Discussion

Our range-wide sampling of *A. lyrata* across Eurasia reveals the extent of *A. lyrata* tetraploids (Figure 1), which was dramatically underestimated until now. Here, we report tetraploids from 30 locations and classify them as autopolyploids, which is apparent from both their deep clustering within *A. lyrata* (Figure 2) and the absence of fixed heterozygosity due to subgenome divergence (Supplementary Figure 3). The autotetraploid populations from Northern Ural and Central Siberia regions group together with local diploids rather than with each other (Figure 2, Supplementary Figure 4b), suggesting different source populations for their origin. Considering that autotetraploid *A. lyrata* populations in Europe also form distinct lineages^33^ and that allotetraploid *A. kamchatica* originated from multiple hybridizations between *A. lyrata* and *A. halleri*^44,65^, we conclude that *A. lyrata* is the most polyploid-rich species complex within *Arabidopsis* genus in both auto– and allopolyploid context. The recurrent formation of tetraploids may imply *A. lyrata* (1) has a propensity for whole genome duplication, perhaps by an increased likelihood of producing unreduced gametes or increased sporadic somatic WGD, (2) establishment of *A. lyrata* tetraploids is facilitated by special environmental conditions or genetic background.

*A. lyrata* is the northernmost species among *Arabidopsis*, and its evolutionary history has been largely affected by temperature fluctuations during the Pleistocene. Expansion of *A. lyrata* eastwards from Europe has been proposed to predate the second to last Ice Age^35^, which peaked at 130 Kya. Pleistocene temperature oscillations and glaciation cycles explain the main migration routes and secondary contacts in refugia areas, such as Beringia^44^. TreeMix analysis grouped Siberian and Karelian populations together, while UK and Scandinavian populations grouped with Central European populations, consistent with previously suggested different colonization routes for Karelia and Scandinavia, with the former being colonized from Northern Ural and the latter from Central Europe, both occurring after the ice sheet retreat post-LGM^48,72^. Such a pattern of distinct re-colonization routes is repeated in *Arabidopsis thaliana*^73^, and other Arctic species including keystone species such as *Dryas octopetala, Vaccinium vitis-idaea* and *Betula nana*^74^.

Given that *A. lyrata* appears to be the most polyploid-rich taxon in the *Arabidopsis* genus, we again ask: what explains the abundance of polyploids in certain lineages, their emergence at certain times and climates? Our estimations of the divergence times between the diploids and tetraploids in Northern Ural and Central Siberia arrive close to the peak of the last Ice Age (Figure 2) and follow a general pattern of co-occurrence with glaciations^25^. This suggests that environment may play a crucial role in polyploid birth rate, triggering whole-genome duplications via production of 2x gametes: either unreduced gametes formed in a diploid, or normal gametes of a somatically doubled branch. Environmental stress can disturb microtubules and prevent the completion of meiotic or mitotic cell division^12,75^.

However, as the emergence of polyploids does not guarantee their survival, what contributes to the reduced death-rate in Siberian tetraploid lineages? While environmental disturbance may also facilitate establishment of polyploids, *e.g.* by opening novel space for colonization after the ice-sheet retreat^76,77^, we see strong evidence of the impact of genetic adaptation. Topology weighting analysis and haplotype sharing from *de novo* assemblies reveal introgression among distinct tetraploid lineages over long distances (Figure 3). Given that the tetraploid populations in Europe were dated at around 160 Kya^39^, considerably older than Siberian tetraploid lineages dated around 16 and 11 Kya, we infer the direction of the gene flow from Central European *A. lyrata* all the way to Central Siberian *A. lyrata* (Figure 3). The interspecific introgression from *A. arenosa* to *A. lyrata* in Central Europe has been shown previously^39^. Regions introgressed from Central Europe to Northern Ural and to Central Siberia are narrow and highly enriched in meiotic genes (Figure 3a,b). Among introgressed haplotypes in Siberia, we find the *ASY3* tetraploid-derived allele which has an established functional role^23^. In natural tetraploid *A. arenosa*, plants with the derived tetraploid *ASY3* allele in conjunction with a derived *ASY1* allele have fewer multivalents and shorter meiotic axes than plants with the diploid alleles^23^, leading to more stable meiosis. Additional research^71^ on *ASY3* as an adaptation candidate showed an increase in meiotic stability associated with the derived tetraploid *ASY3* haplotype. During meiosis, the length of the chromosomal axis dictates crossover frequency and position^78,79^. In tetraploids, axis length may be especially important if it serves to limit crossovers to one per chromosome, reducing the risk of multivalent formation^80^. Patterns of diversity and divergence in Siberian lineages at the introgressed region harboring *ASY3* gene are consistant with positive selection, suggesting that introgression was adaptive.

Similar patterns of both long-range introgression and positive selection in *A. lyrata* polyploids across Eurasia are observed in the region harboring the cohesin cofactor PDS5. In budding yeast, PDS5 supports sister chromatid cohesion, as mutants suffer from early separation of sister chromatids^81^. While experimental work in *A. thaliana* shows that PDS5 mutants have minimal meiotic phenotypes^82^, suggesting its role in *Arabidopsis* diverges from that in yeast systems, more recent work shows that PDS5 impacts axis length. As described above for ASY3, regulation of axis length in tetraploids can help to stabilize meiosis, as a shorter axis can limit crossover number. *PDS5* and its paralogs were repeatedly found in selection scans between diploid and tetraploid lineages in Brassicaceae: first in *A. arenosa*^20,83,84^, then in Central European *A. lyrata*^39^, and in *Cardamine amara*^85^, where tetraploids harbor a private derived allele, absent from diploid gene pools. We find a high frequency of described adaptive tetraploid alleles in *ASY3* and *PDS5* meiotic genes in Northern Ural and Central Siberian tetraploids, which suggests that meiotic stabilization is required for polyploid establishment, and that this stabilization is primarily achieved through adaptive introgression.

Siberian *A. lyrata* tetraploids are younger compared to European and it is possible that meiotic stabilization there is incomplete. For example, we find synaptonemal complex protein ZYP1b in introgression scans across Eurasia. The paralogs *ZYP1a* and *ZYP1b* encode transverse filament proteins which are central elements of the synaptonemal complex, helping to “zip” together chromosomal axes of homologs, distribute cross-overs and establish proper bivalents^86–88^. Tetraploid alleles of *ZYP1* paralogs are likely adaptive as they appear on introgression and selection scans^20,39,84^. However, in Siberia, although tetraploid Central European alleles of *ZYP1b* are already introduced and detected by introgression scans, their frequency is still low as *ZYP1b* does not appear among outliers in diversity and divergence patterns (Figure 3). Meiotic adaptation to polyploidy is polygenic without a clear key player but rather many players of cumulative effects^17,89^, the dynamics of which we probably observe here.

It seems that, while environmental fluctuation may yield increased polyploid formation in *A. lyrata*, the persistence of these neopolyploids is due to genetic adaptation involving long-range introgression. This raises the question of how neopolyploids survived at first. In contrast to *A. arenosa*, *A. lyrata* is known to propagate clonally (via root sprouts), which could help local neopolyploid persistence in spite of initial unstable meiosis. Environmental conditions could also play a role in early stabilization. For example, it is known that temperature impacts meiosis by modifying the number of crossovers per chromosome^90^. It is plausible that vegetative propagation and environmental conditions allowed the initial survival of neopolyploid *A. lyrata* in Siberia. Then, gene flow into the nascent tetraploid populations from the established Central European gene pool introduced the adaptive alleles, enhancing further polyploid establishment and ultimately resulting in further persistence of polyploids in Siberia.

## Materials and Methods

### Fieldwork and plant collection

We used a combination of herbarium specimens and seeds collected from the field for this study. For herbarium accessions, we sampled a small amount of dried tissue from at least one individual plant on a herbarium sheet. Samples from a single collection locale were named using the herbarium accession appended by a number indicating multiple plants that were sequenced from a sheet (e.g. MW0079552_1, MW0079552_2 and MW0079552_3). We collected seeds from wild populations of *A. lyrata* in expeditions to the Lena river valley, Taz Estuary, Polar Urals and South Yakutia (Supplementary Data 1). We then grew these seeds in the greenhouse (21 C°, 16 hours of daylight) and sampled leaf tissue for sequencing.

### Short-read sequencing for population analyses

Plant material was processed in two different ways, indicated by types I and II in Supplementary Data 1. Processing protocols for each type are described in detail in ^62^. Briefly, we isolated DNA from type I herbarium samples using the DNeasy Plant Mini Kit (Qiagen) followed by library prep with the NEBNext® DNA Library Prep Kit. For type II samples, we isolated DNA using the NucleoMag© Plant” kit from Macherey and Nagel (Düren, Germany) on the KingFisher 96Plex device (Thermo), followed by a TPase-based library prep. Libraries were pooled and sequenced on the NovaSeq 6000 S4 flow cell Illumina system. Full list of short-read sequenced samples used in the study is in the Supplementary Data 1.

### Linked-read sequencing

DNA from 2 g of fresh material was isolated with a NucleoBond HMW DNA kit (Macherey Nagel). Quality, library preparation and sequencing was performed by BGI as described in ^91^. Library preparation process starts from random insertions of a transposon sequence into the long fragment DNA, then the product is combined with a magnetic bead carrying a multi-copy molecular barcode. Each DNA molecule receives a unique barcode. After that fragments are amplified using PCR, and single-stranded cyclized products are used for pair-end sequencing DNBSEQ with read length of 100bp.

### Short read mapping and variant calling for population analysis

We analyzed short paired-end reads (2 x 150 bp) the same way as in Kolesnikova et al^62^. The reads were filtered for adapters with bbduk.sh script from BBMap (38.20)^92^ with ktrim=r k=23 mink=11 hdist=1 tbo tpe qtrim=rl trimq=15 minlen=70. Then, we mapped reads to the NT1 *A. lyrata* reference genome using bwa mem (0.7.17)^93^ with –M parameter. We used picard MarkDuplicates^94^ to mark PCR duplicated reads and then used samtools^95^ to sort and index the bam files. We called variants with HaplotypeCaller algorithm from GATK (3.8)^96^ and then combined individual vcf files using CombineGVCFs and created final vcf using GenotypeGVCF from GATK including non-variant sites with –-includeNonVariantSites parameter. We estimated heterozygosity on a vcf with variants called for all samples as for diploids by calculating proportion of heterozygous sites among all called sites. For the further analysis we created a vcf with variant and non-variant sites called according to the ploidy of the samples (HaplotypeCaller with –-sample-ploidy 2 or –-sample-ploidy 4).

### Synthetic long-read assemblies and tree inference

StLFR reads were assembled using the stlfr2supernova_pipeline (https://github.com/BGI-Qingdao/stlfr2supernova_pipeline) which is based on Supernova^97^ with –-style=pseudohap2 option to get two haplotypes for each assembly. The haplotypes were scaffolded on the NT1 *A.lyrata* genome^62^ with ragtag^98^. GALBA pipeline^99–103^ was used for protein annotation of the assemblies with proteins from Phytozome^104^ *A.lyrata* v.2.1 release^105^ as a protein database. The annotated genes were used for gene tree construction. Protein BLAST^106^ was used to identify genes of interest in the annotation. Then, we used L-INS-i algorithm of MAFFT^107^ for multiple sequence alignment of the gene regions. The alignments were manually curated and used for tree construction with iqtree^108,109^, fast bootstrap=1000.

### Ploidy estimation and *S*-allele genotyping from short-read sequencing data

To estimate ploidy we used nQuire^53^ with “create”, “denoise” and “lrdmodel” steps. This ploidy estimation method is based on allelic read depth distributions, also previously described^110^. We identified *S*-alleles as described in Kolesnikova et al^62^ using short-read sequencing data and *S*-locus genotyping pipeline NGSgenotyp^54^ with two first steps: kmerfilter (-k 20 for *SRK* and –k 15 for *SCR*) and genotyp – with both *SRK* and *SCR* sequence reference databases. In addition to the databases used previously^62^, we have assembled four *SRK* alleles from short-read data with assembly option and provide them together with 6 more published^59,111^ *SRK* alleles in Supplementary Table 3. The sequences of *S*-alleles are provided in the Supplementary Data 3 in .fasta format. The number of different *S*-alleles in each individual together with proportion of heterozygous sites were used to confirm the estimated ploidy level.

### Visualization

We used R v4.2.2^112^, primarily with the tidyverse v2.0.0 package collection^113^, for making the plots. Maps were created using Lambert Asimuthal Equal Area projection centered at N 40° E 90°, packages sf v1.0-14^114^ and raster v3.6-23^115^ and a background map by NaturalEarth; the ggrepel algorithm^116^ was used to adjust the positioning of the pie charts. The Last Glacial Maximum (LGM) ice sheet extents^69^ were simplified for plotting.

### Cytogenetic validation of ploidy

To confirm ploidy levels estimated by bioinformatic methods, we used meiotic spreads. We first harvested immature buds (prior to pollen being formed) and fixed them in Carnoy (3:1; Ethanol: Acetic Acid) for 2h on a horizontal shaker at room temperature, changed to fresh fixative, and stored them at –20°C until further usage.

Fixed buds were rinsed twice for 5min in 10mM Citrate buffer (tri-sodium citrate, pH 4,5 with HCl), followed by a digestion with an enzymatic solution (0,3% Pectolyase Y-23, 0,3% Driselase, 0,3% Cellulase Onozuka R 10 and 0,1% Sodium Azide in 10mM Citrate buffer) at 37°C for 1h in a humid chamber. After another two washes with Citrate buffer, the anthers were separated from surrounding tissue and put onto a slide. One edge of an anther was cut open and the cells within were squeezed out with the tip of a needle. After processing all anthers of 1-2 buds, cells were mixed into a suspension by stirring with a needle. A small amount of freshly prepared 60% Acetic Acid was added to the suspension from the side and the whole drop gently homogenized by again stirring with a needle. The slide was placed on a hot plate at 45°C and cells were spread with the help of a bent needle until almost all liquid evaporated. Around the spread area, a boundary of ice cold fresh fixative solution was created and allowed to enter, followed by a jet of ice cold fixative directly into the center of the area. Then the slide was allowed to air dry in an upright position. Chromosomal Spreads were stained with DAPI in a solution of 2ug/ml in the antifade mounting medium Vectashield and covered with a coverslip. Imaging was done on a Zeiss Axio Imager Z2 Microscope using the ZEN software. Ploidy validation data shown in Supplementary Data 2.

### Estimation of nucleotide diversity

We calculated π^117^ for diploid and tetraploid live populations with more than five individuals in each (excluding siblings) using piawka (https://github.com/novikovalab/piawka), our implementation of the site weighting strategy introduced in pixy^118^ compatible with mixed-ploidy datasets. We also calculated π within each tetraploid lineage and between diploid-tetraploid lineages on a per-gene basis using piawka.

### Twisst

Starting with a VCF containing only biallelic SNPs, we filtered the dataset further using genomics_general-0.4 toolkit (minimum quality=20, minimum depth=5, maximum missing data=0.2, maximum heterozygosity=0.75, minimum variant count=5). Genealogies were inferred in windows of 50 SNPs along the genome using PHYML^119^ and a phasing approach implemented in genomics_general. Previously published^45^ samples of *Arabidopsis cebennensis* and *Arabidopsis pedemontana* were used as outgroups. We used Twisst^68^ for topology weighting. GO enrichment analysis using g:Profiler was performed for genes intersecting the regions with >50% weighting of introgression topologies.

### Demographic Modeling

To generate Site Frequency Spectra (SFS), we utilized biallelic SNPs and invariant sites. We excluded genic, centromeric and pericentromeric regions, retaining only SNPs found in intergenic regions of chromosome arms. Individuals of the two main lineages were selected as outlined in Supplementary Data 1. Using the chosen SNPs we generated bootstrap replicates by dividing the genome into 10kb segments. These segments were resampled with replacement until the equivalent size of the total amount of filtered SNPs in the genome. We computed site frequency spectrum (SFS) for each population of the 200 boot-strapped replicates and a joint site frequency spectrum for each pseudoobservation using the python package dadi^120^. The SFS spectra generated by dadi were used as input into fastsimcoal2 (fsc27)^70^. We first assessed a simple divergence model for both lineages through 50 fastsimcoal2 runs, then compared with a model allowing migration from the diploid into tetraploid lineage. Each fastsimcoal run was allowed 40 optimization cycles and 100,000 iterations. Based on likelihood and AIC criteria we continued with the divergence mode with migration for the confidence interval calculation. For each of the 200 bootstrap replicates we initiated 50 fastsimcoal2 runs. The resulting parameter estimates from the 200 pseudo-observations were used to calculate the 95% Confidence Intervals (CIs) in R.

### Data availability

The whole genome re-sequencing short reads data for the samples which was generated in this study were submitted to the ENA database under study number PRJEB67879 and PRJEB60410. The list of all the samples used in the study with individual ENA numbers is in Supplementary Data 1. The images of the herbarium vouchers can be found online using an example link https://plant.depo.msu.ru/open/public/item/MW0079552 and replacing MW0079552 with the individual number from Supplementary Data 1 column “accession”. Both raw stLFR data and resulting assemblies are also accessible in the ENA database, with individual ENA numbers in Supplementary Data 5. Supplementary Data 1-5 and the scripts used in the analysis are available at https://github.com/novikovalab/Siberian_Alyrata

## Acknowledgments

We thank the numerous collectors of the herbarium material we used and the herbarium curators A.V. Verkhozina (IRK) and N.V. Stepantsova (IRKU) who kindly provided access to some collections. We are grateful to Igor Pospelov (FGBU “Zapovedniki Taimyra”) and Elena Pospelova for the live samples from the Putorana massif and to Daiana Zhernova for the live samples from the White Sea coast. Our fieldwork was additionally supported by the Austrian Science Fund (FWF grant P30208-B-29) and held within the state assignment of the Papanin Institute for Biology of Inland Waters Russian Academy of Sciences (theme 121051100099-5). We also owe its success to many locals and other friends who provided us shelter and transport. Special thanks to Tatiana Kirillina, Matryona Achikasova-Koryakina, Djulustan Nikitin and Chekurovka camp fishermen, Aleksandr Markevich, Manchaary Vladimirov, Stolb weather station staff, Khokhuccha camp fishermen, staff of Zhigansk and Bykovsky schools, Leonid Shaglanov, Tiksi Arctic Gymnasium staff, Grigorii from Sagastyr, Aleksandr Gukov, Andrey Kuzkin and Yevgeny Serebrennikov (Lena river area, 2019–2020); Vladimir Segoy together with Samburg firefighters, Prokopiy Tesida, staff of the Messo-Yakha protected area, Aleksey Yar and nenets communities of Salyakaptan, Salyapayuta and Yabtasalya areas (West Siberia, 2021); and Faina and her family from the Kharp town (Polar Urals, 2021). The work of A.P.S. on the curation of dry plant material was supported by the Russian Science Foundation (project #21-77-20042). We thank the Max Planck-Genome-Centre Cologne, Germany (http://mpgc.mpipz.mpg.de/home/) for performing short-read libraries in this study. We thank Levi Yant and Andrew C. Clarke for performing DNA extractions and sequencing of a subset of the herbarium samples. The sequencing and data analysis were funded by the European Union (ERC, HOW2DOUBLE, 101041354) and by the joint Deutsche Forschungsgemeinschaft (DFG) and GACR program —project number 490698526. This research was also funded by Czech Science Foundation (project 22-29078K) to FK. The views and opinions expressed are however those of the author(s) only and do not necessarily reflect those of the European Union or the European Research Council Executive Agency. Neither the European Union nor the granting authority can be held responsible for them.

