## Supplemental Materials for "Multiple polyploidizations in *Arabidopsis lyrata* stabilized by long-range adaptive introgression across Eurasia"

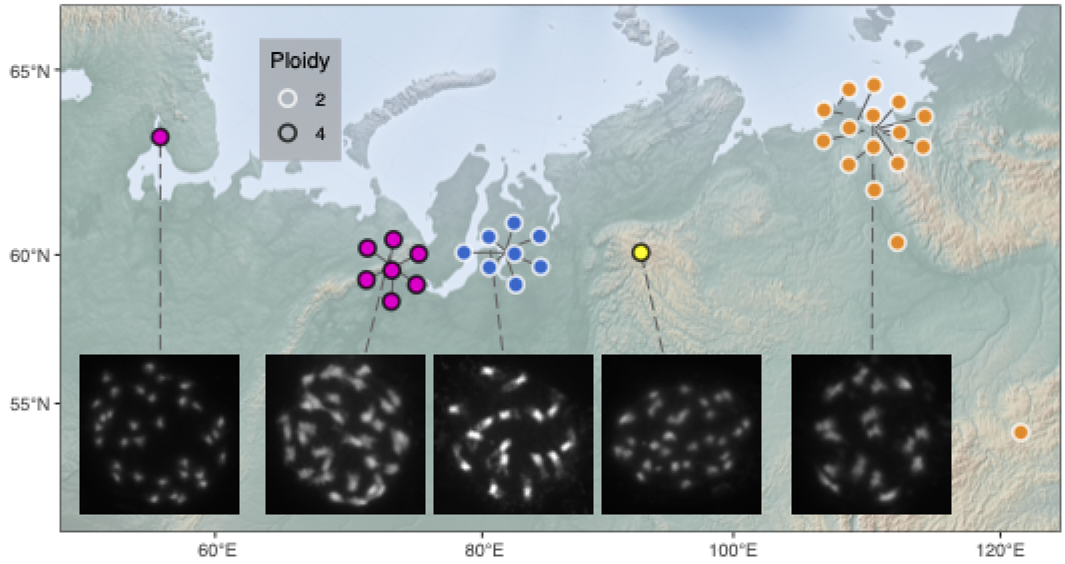

Supplementary Figure 1. Geographical map of populations with collected seeds. Seeds were grown in the green house and karyotyped to confirm the ploidy inference of each lineage on Figure 2.

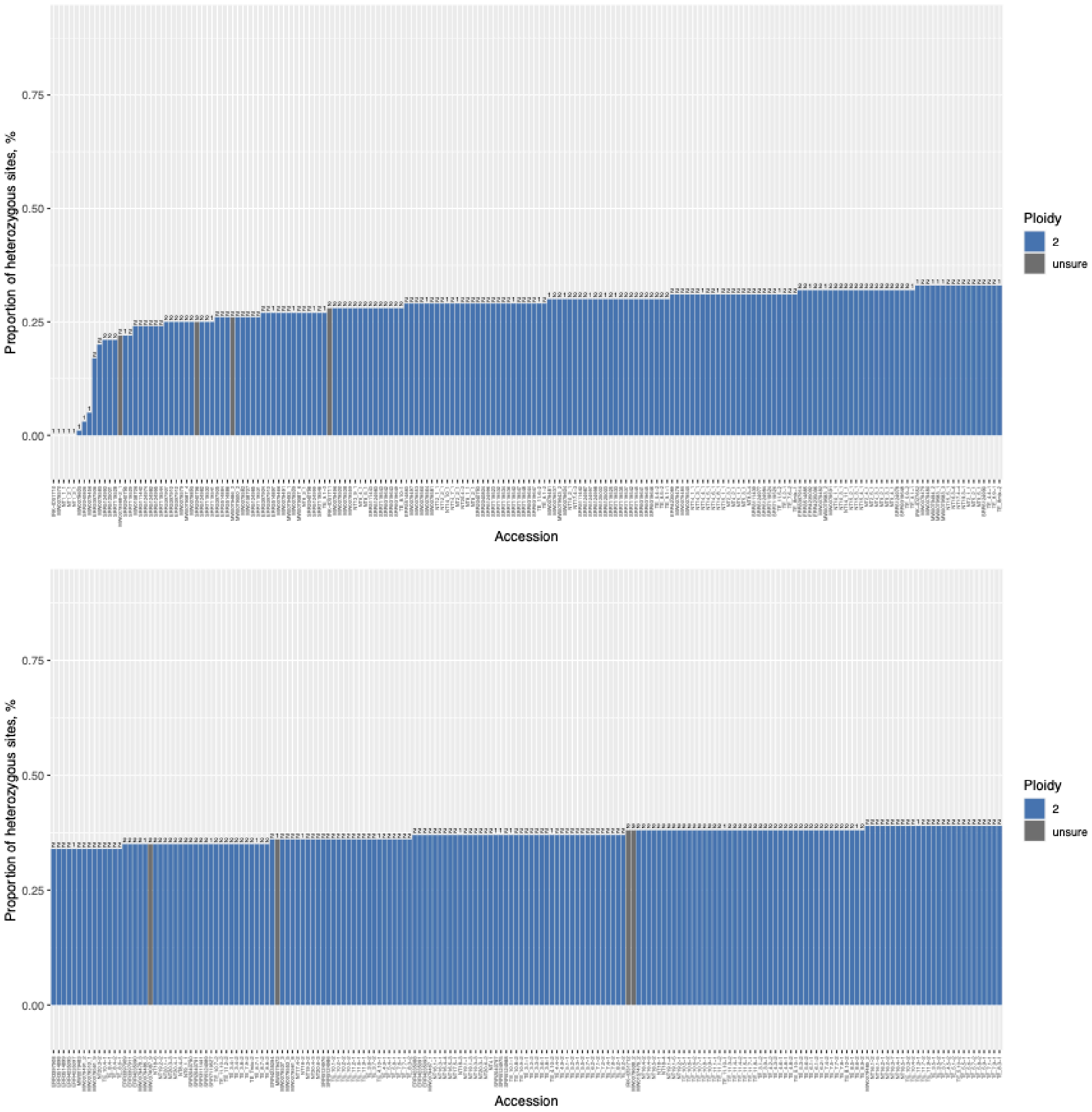

Proportion of heterozygous sites, %
0.75
0.50
0.25
0.00
Accession
Ploidy
2
unsure
Proportion of heterozygous sites, %
0.75
0.50
0.25
0.00
Accession
Ploidy
2
unsure

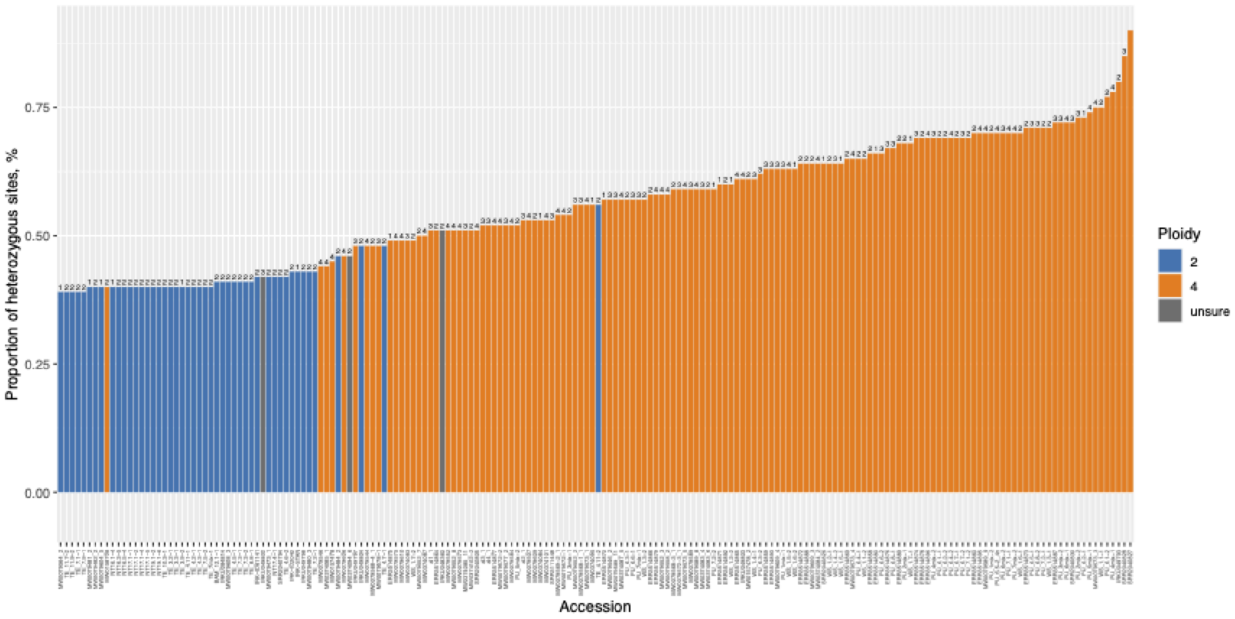

Supplementary Figure 2. Proportion of heterozygous sites corresponds to inferred ploidy by nQuire, marked by color, and number of different *S*-alleles indicated on top of each bar.

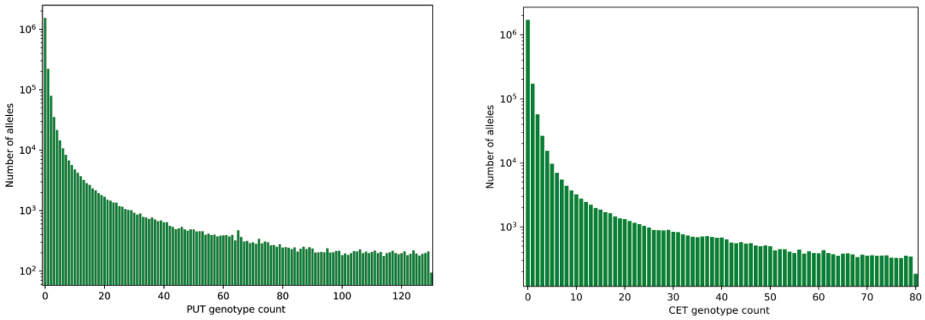

Supplementary Figure 3. Site frequency spectra of the Northern Ural and Central Siberia tetraploids, showing no peak at the intermediate frequencies

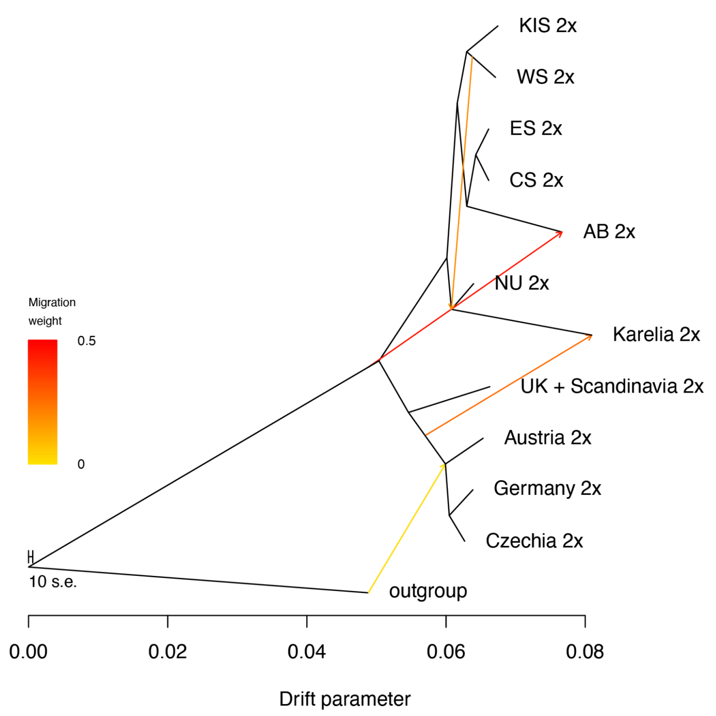
KIS 2x
WS 2x
ES 2x
CS 2x
AB 2x
NU 2x
Karelia 2x
UK + Scandinavia 2x
Austria 2x
Germany 2x
Czechia 2x
outgroup
Migration weight
0.5
0
10 s.e.
0.00
0.02
0.04
0.06
0.08
Drift parameter

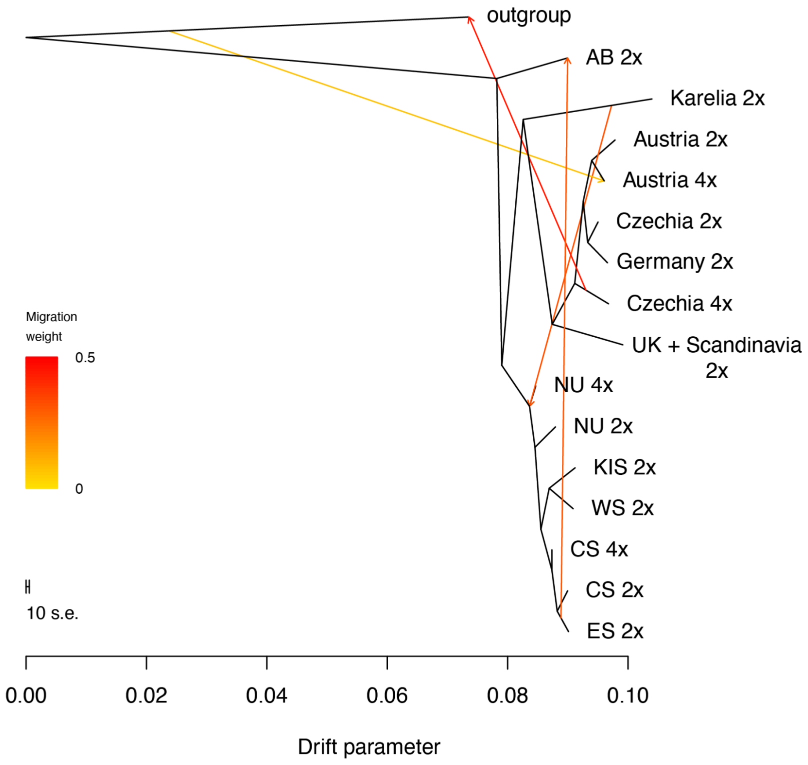

Supplementary Figure 4 Maximum likelihood TreeMix models including diploids only (top) and with tetraploids (bottom). NU - Northern Ural, WS - Western Siberia, CS - Central Siberia, ES - Eastern Siberia, KIS - Kuz’kin Island, AB - Amur Basin.

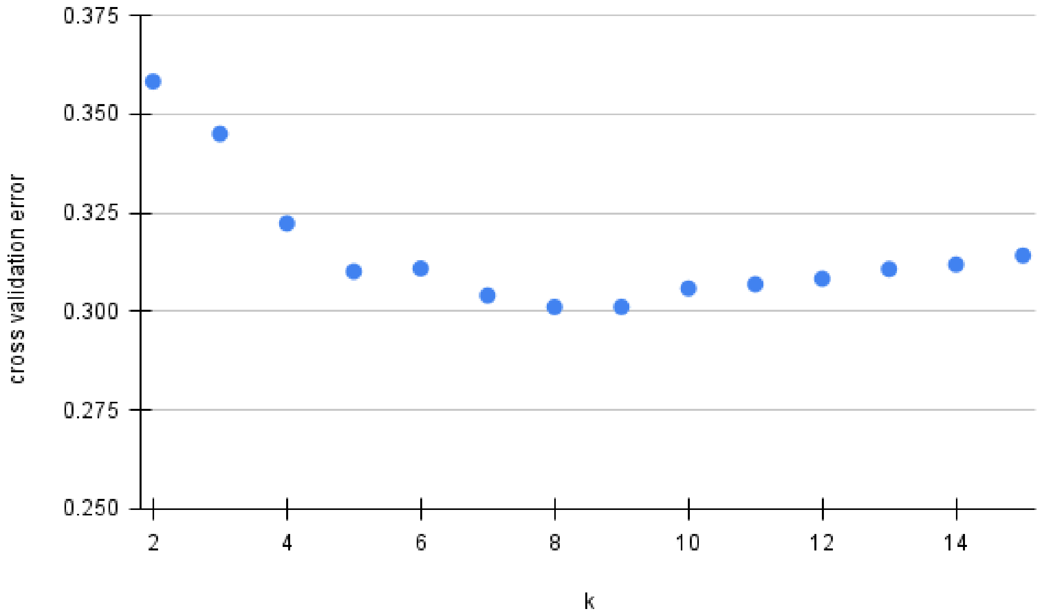

Supplementary Figure 5. Cross-validation error plot to choose k for admixture. K=8 was chosen as an optimal number of clusters for Figure 2a

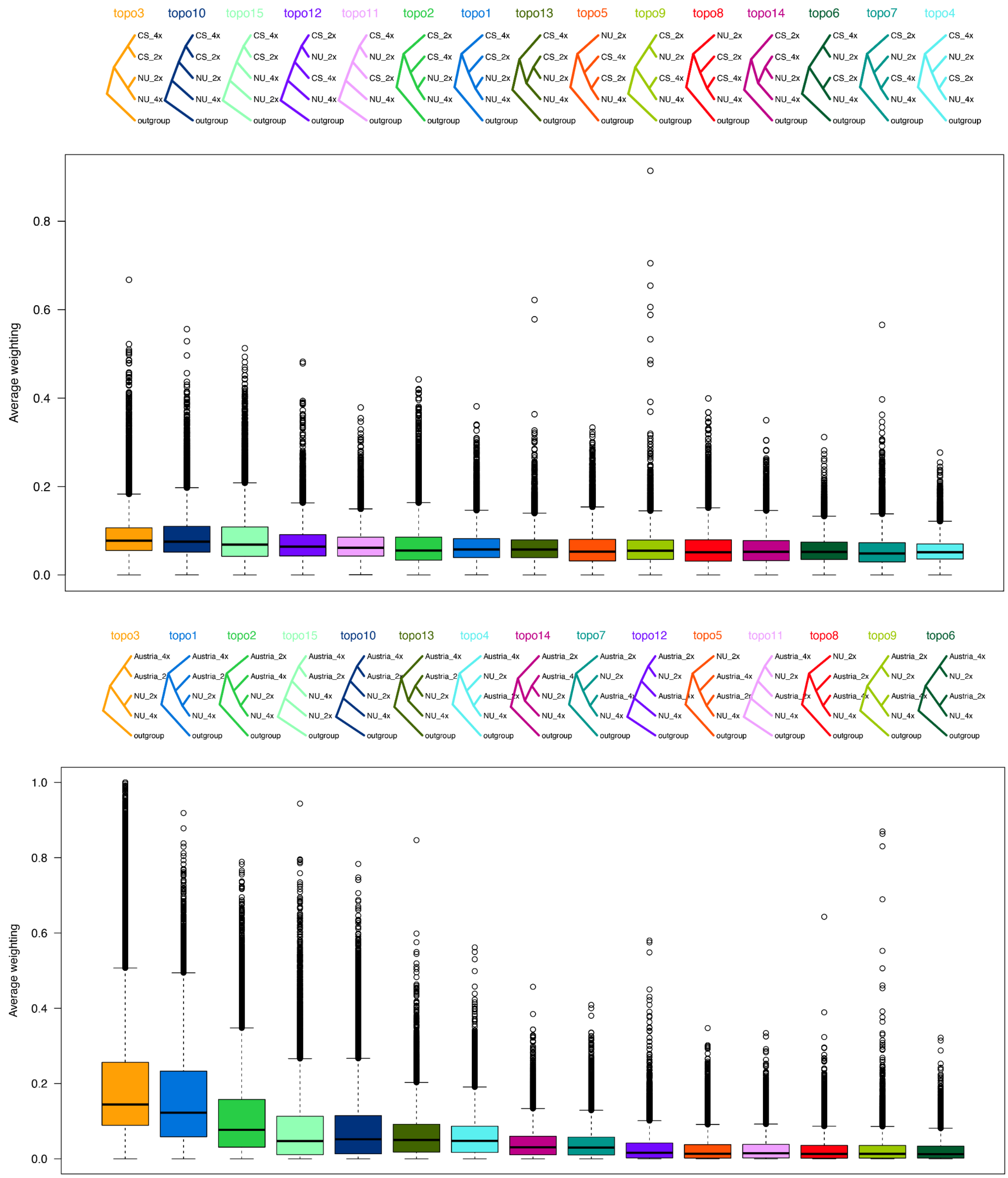

Supplementary Figure 6. Full topology weighting results across all possible topologies. Upper plot, topology weighting for Northern Ural and Central Siberian lineages. Lower plot, topology weighting for Northern Ural and Central Europe lineages.

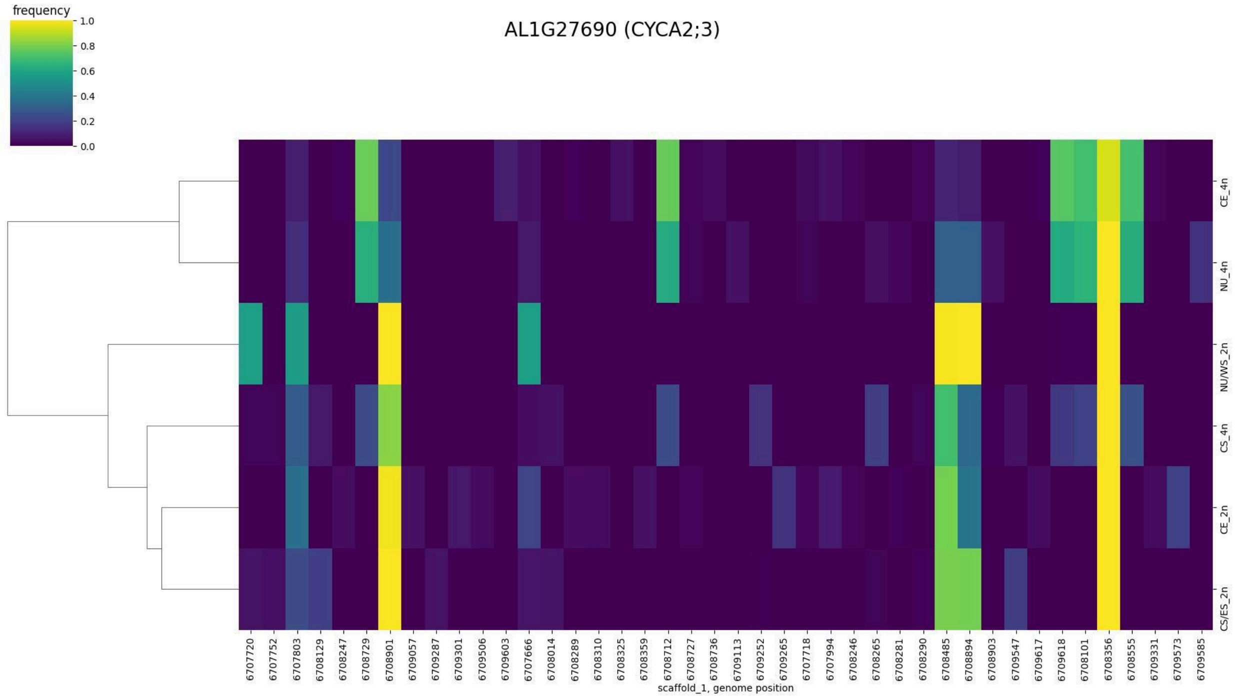
AL1G27690 (CYCA2;3)
frequency
1.0
0.8
0.6
0.4
0.2
0.0
CE_4n
NU_4n
NU/WS_2n
CS_4n
CE_2n
CS/ES_2n
6707720 6707752 6707803 6708129 6708247 6708729 6708901 6709057 6709287 6709301 6709506 6709603 6707666 6708014 6708289 6708310 6708325 6708359 6708712 6708727 6708736 6709113 6709252 6709265 6707718 6707994 6708246 6708265 6708281 6708290 6708485 6708894 6708903 6709547 6709617 6709618 6708101 6708356 6708555 6709331 6709573 6709585
scaffold_1, genome position

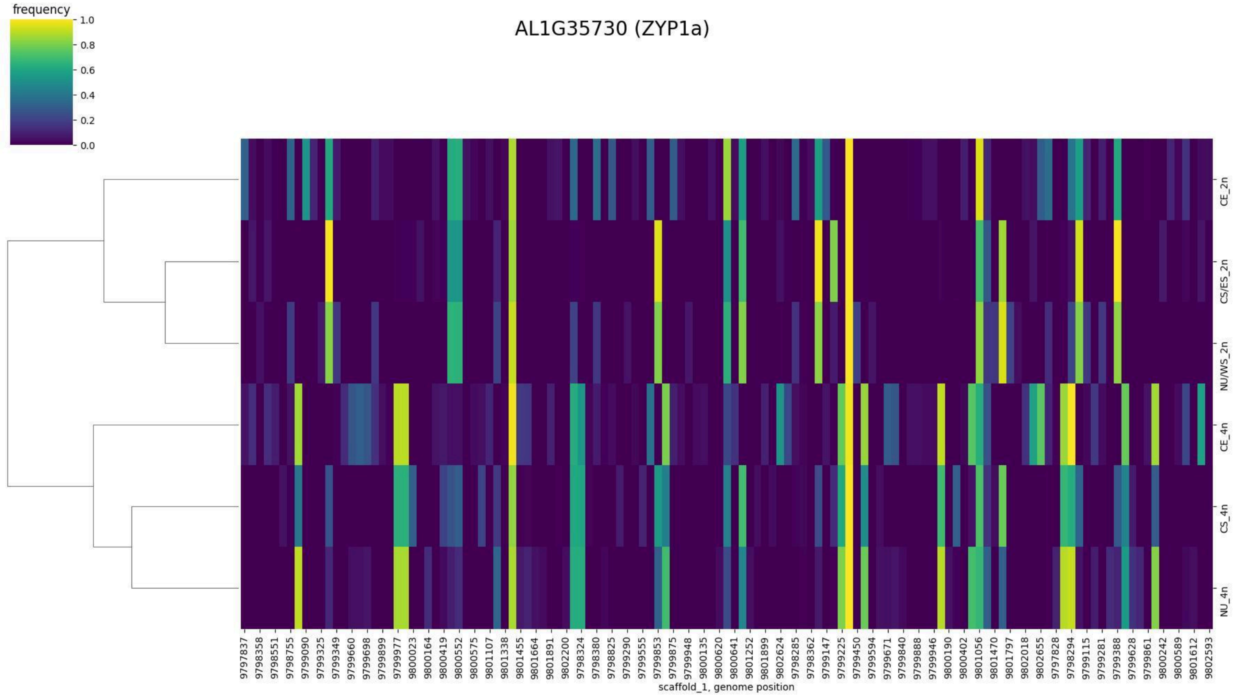
AL1G35730 (ZYP1a)
frequency
1.0
0.8
0.6
0.4
0.2
0.0
CE_2n
CS/ES_2n
NU/WS_2n
CE_4n
CS_4n
NU_4n
9797837 9798358 9798551 9798755 9799090 9799325 9799349 9799660 9799698 9799899 9799977 9800023 9800164 9800419 9800552 9800575 9801107 9801338 9801455 9801664 9801891 9802200 9798324 9798380 9798825 9799290 9799555 9799853 9799875 9799948 9800135 9800620 9800641 9801252 9801899 9802624 9798285 9798362 9799147 9799225 9799450 9799594 9799671 9799840 9799888 9799946 9800190 9800402 9801056 9801470 9801797 9802018 9802655 9797828 9798294 9799115 9799281 9799388 9799628 9799861 9800242 9800589 9801612 9802593
scaffold_1, genome position

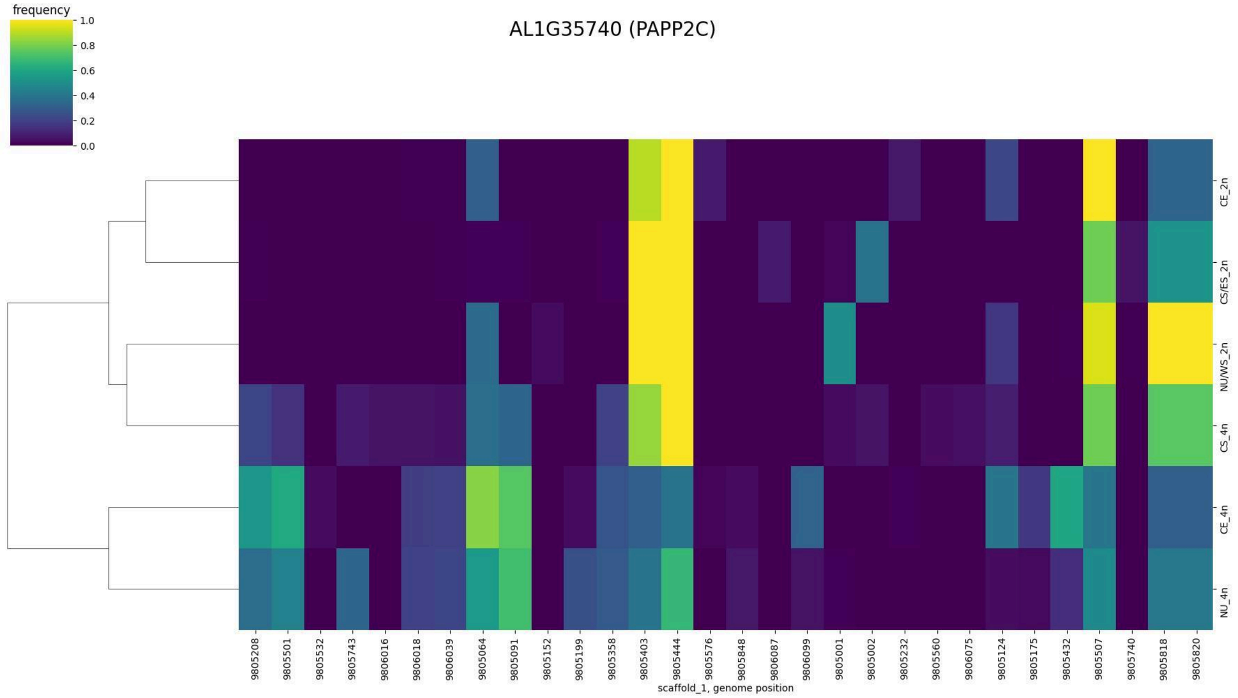
AL1G35740 (PAPP2C)
frequency
1.0
0.8
0.6
0.4
0.2
0.0
CE_2n
CS/ES_2n
NU/WS_2n
CS_4n
CE_4n
NU_4n
9805208
9805501
9805532
9805743
9806016
9806018
9806039
9805064
9805091
9805152
9805199
9805358
9805403
9805444
9805576
9805848
9806087
9806099
9805001
9805002
9805232
9805560
9806075
9805124
9805175
9805432
9805507
9805740
9805818
9805820
scaffold_1, genome position

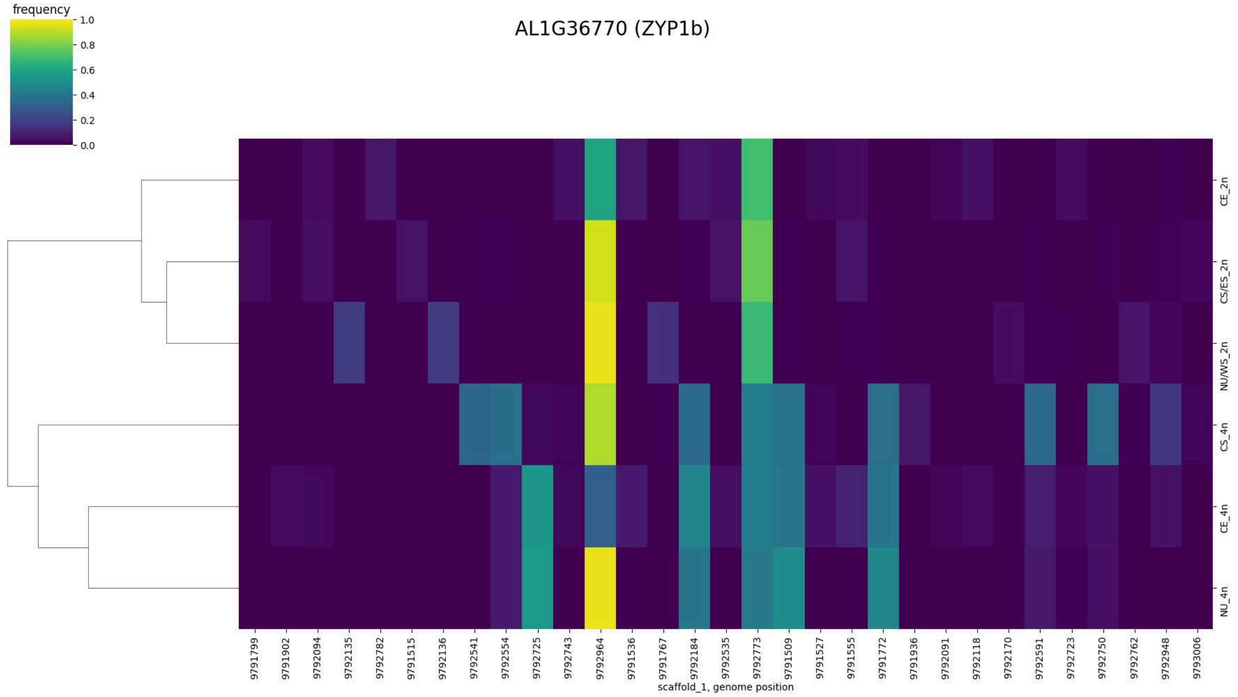
AL1G36770 (ZYP1b)
frequency
1.0
0.8
0.6
0.4
0.2
0.0
CE_2n
CS/ES_2n
NU/WS_2n
CS_4n
CE_4n
NU_4n
9791799
9791902
9792094
9792135
9792782
9791515
9792136
9792541
9792554
9792725
9792743
9792964
9791536
9791767
9792184
9792535
9792773
9791509
9791527
9791555
9791772
9791936
9792091
9792118
9792170
9792591
9792723
9792750
9792762
9792948
9793006
scaffold_1, genome position

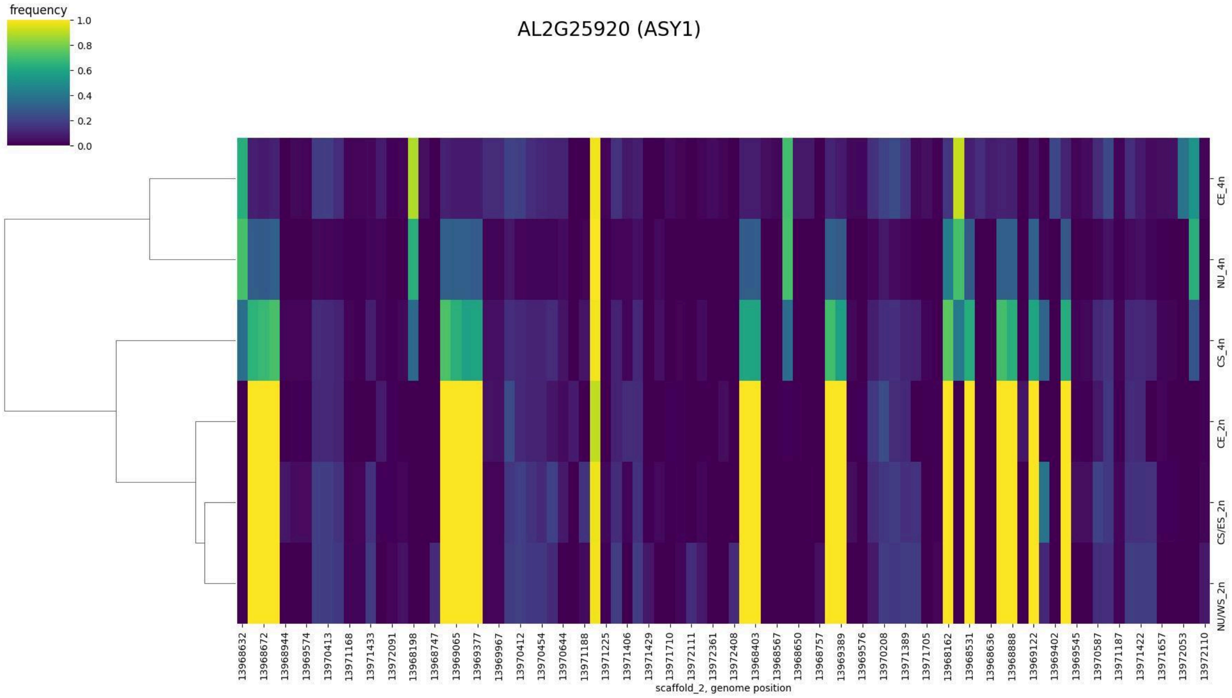
frequency
1.0
0.8
0.6
0.4
0.2
0.0
AL2G25920 (ASY1)
CE_4n
NU_4n
CS_4n
CE_2n
CS/ES_2n
NU/WS_2n
13968632
13968672
13968944
13969574
13970413
13971168
13971433
13972091
13968198
13968747
13969065
13969377
13969967
13970412
13970454
13970644
13971188
13971225
13971406
13971429
13971710
13972111
13972361
13972408
13968403
13968567
13968650
13968757
13969389
13969576
13970208
13971389
13971705
13968162
13968531
13968636
13968888
13969122
13969402
13969545
13970587
13971187
13971422
13971657
13972053
13972110
scaffold_2, genome position

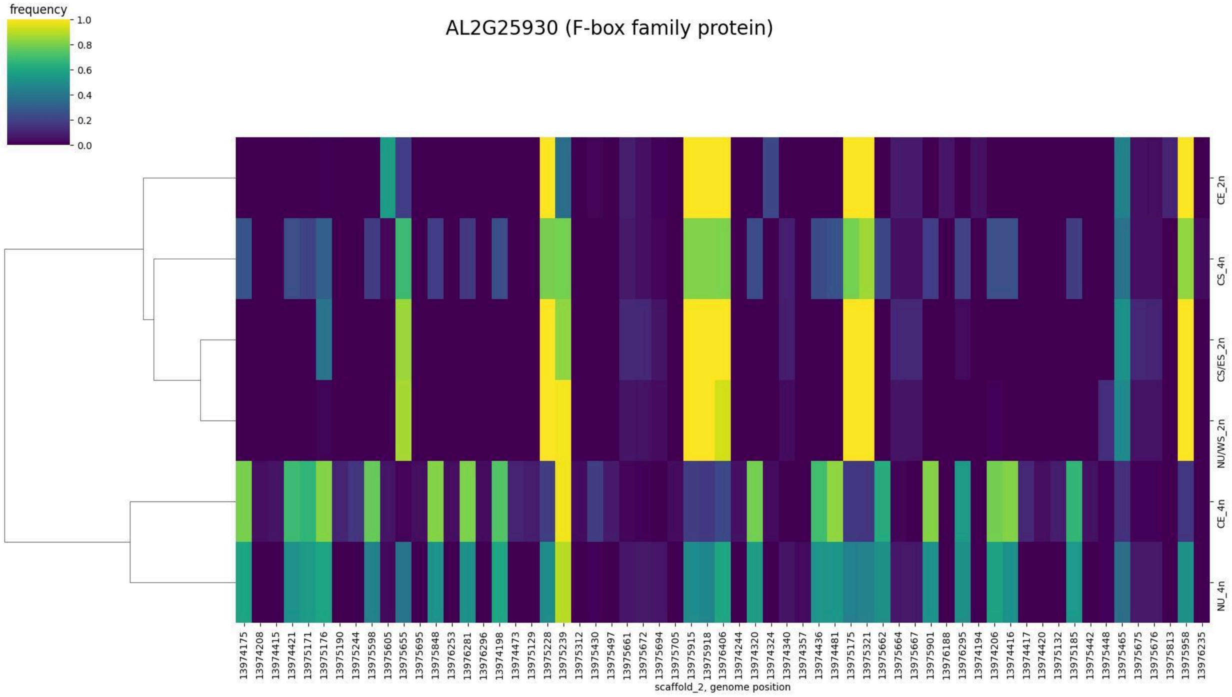
frequency
1.0
0.8
0.6
0.4
0.2
0.0
AL2G25930 (F-box family protein)
CE_2n
CS_4n
CS/ES_2n
NU/WS_2n
CE_4n
NU_4n
13974175
13974208
13974415
13974421
13975171
13975176
13975190
13975244
13975598
13975605
13975655
13975695
13975848
13976253
13976281
13976296
13974198
13974473
13975129
13975228
13975239
13975312
13975430
13975497
13975661
13975672
13975694
13975705
13975915
13975918
13976406
13974244
13974320
13974324
13974340
13974357
13974436
13974481
13975175
13975321
13975662
13975664
13975667
13975901
13976188
13976295
13974194
13974206
13974416
13974417
13974420
13975132
13975185
13975442
13975448
13975465
13975675
13975676
13975813
13975958
13976235
scaffold_2, genome position

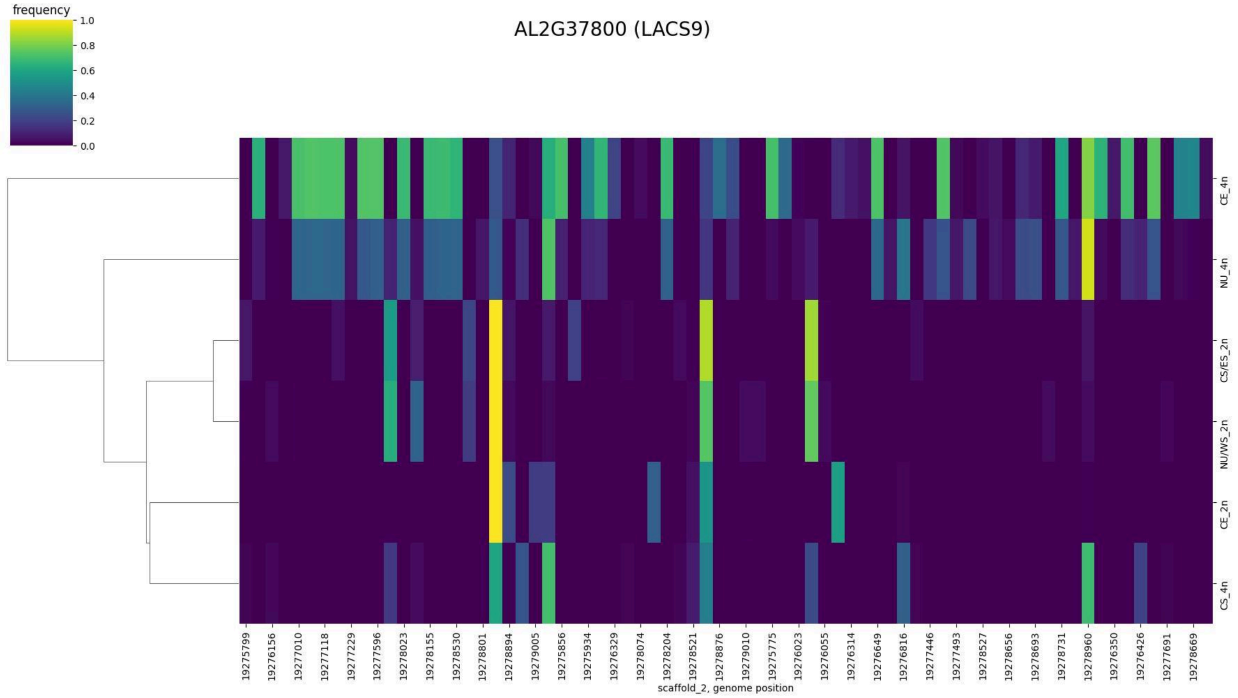
AL2G37800 (LACS9)
frequency
scaffold_2, genome position
CE_4n
NU_4n
CS/ES_2n
NU/WS_2n
CE_2n
CS_4n

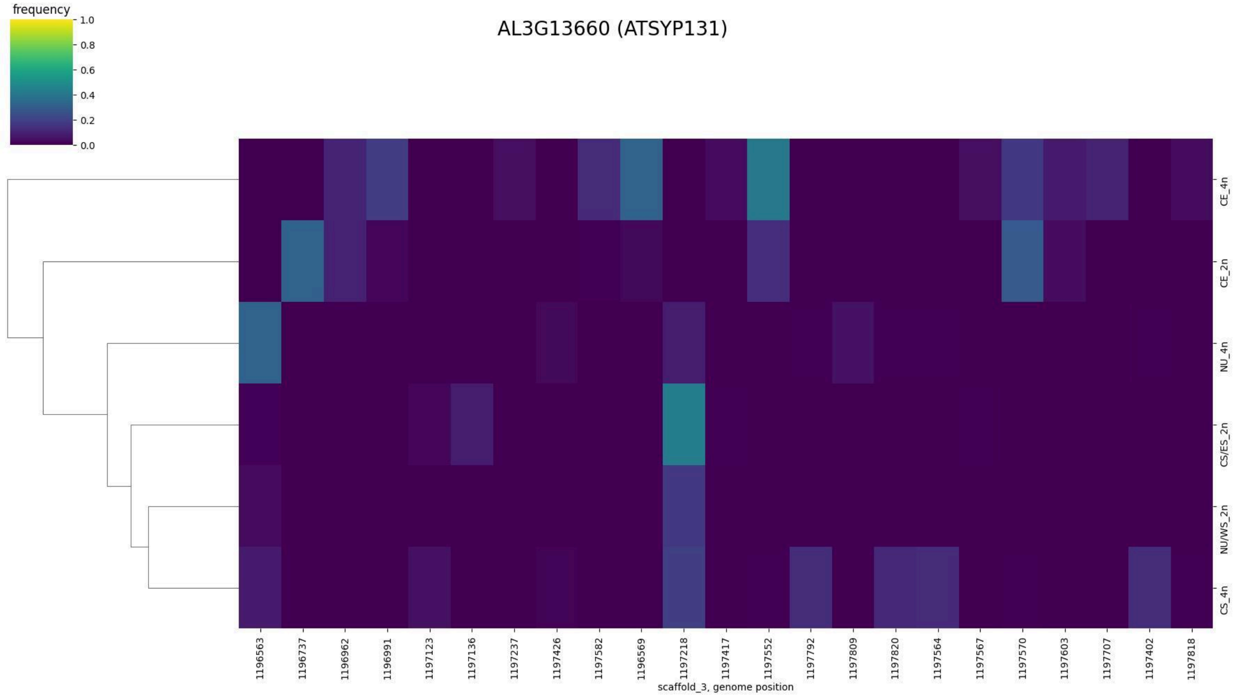
AL3G13660 (ATSYP131)
frequency
scaffold_3, genome position
CE_4n
CE_2n
NU_4n
CS/ES_2n
NU/WS_2n
CS_4n

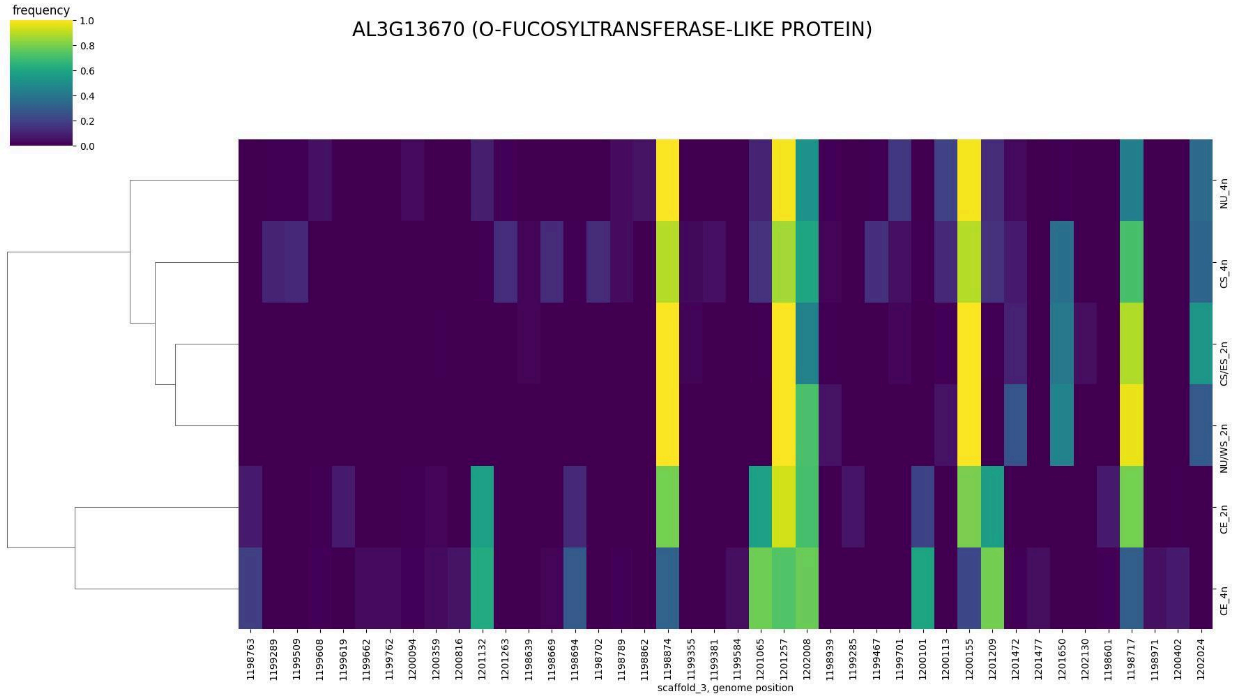
AL3G13670 (O-FUCOSYLTRANSFERASE-LIKE PROTEIN)
frequency
1.0
0.8
0.6
0.4
0.2
0.0
NU_4n
CS_4n
CS/ES_2n
NU/WS_2n
CE_2n
CE_4n
1198763
1199289
1199509
1199608
1199619
1199662
1199762
1200094
1200359
1200816
1201132
1201263
1198639
1198669
1198694
1198702
1198789
1198862
1198874
1199355
1199381
1199584
1201065
1201257
1202008
1198939
1199285
1199467
1199701
1200101
1200113
1200155
1201209
1201472
1201477
1201650
1202130
1198601
1198717
1198971
1200402
1202024
scaffold_3, genome position

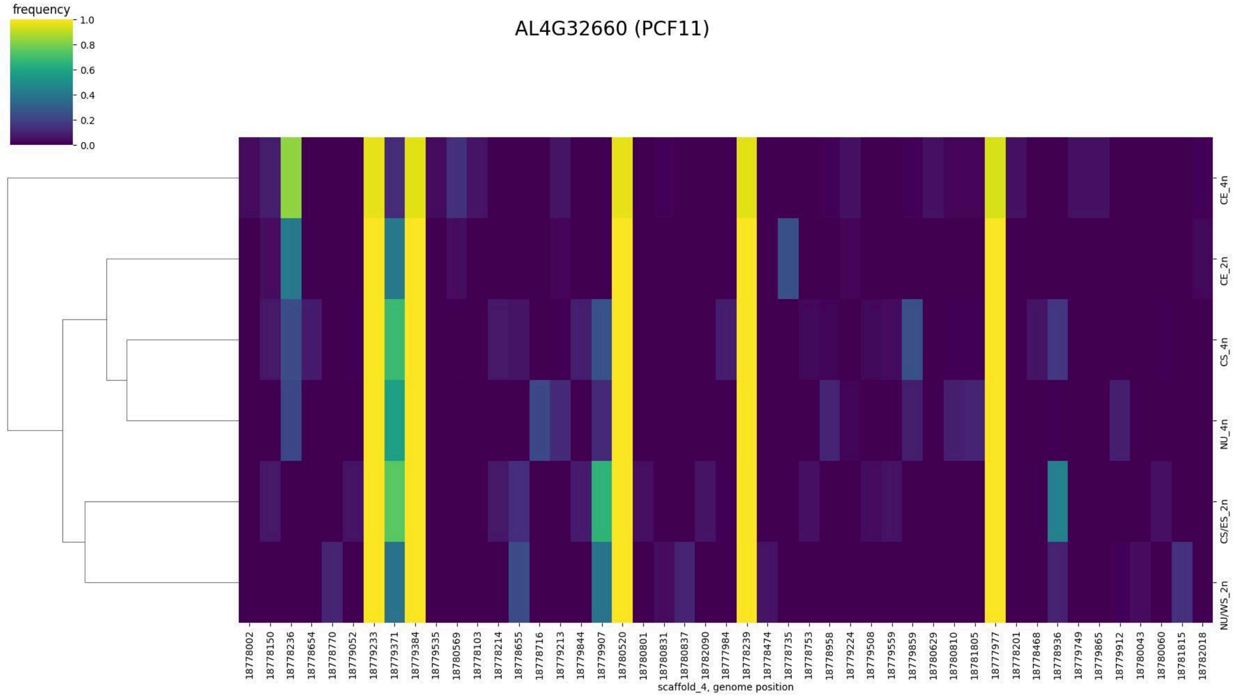
AL4G32660 (PCF11)
frequency
1.0
0.8
0.6
0.4
0.2
0.0
CE_4n
CE_2n
CS_4n
NU_4n
CS/ES_2n
NU/WS_2n
18778002
18778150
18778236
18778654
18778770
18779052
18779233
18779371
18779384
18779535
18780569
18778103
18778214
18778655
18778716
18779213
18779844
18779907
18780520
18780801
18780831
18780837
18782090
18777984
18778239
18778474
18778735
18778753
18778958
18779224
18779508
18779559
18779859
18780629
18780810
18781805
18777977
18778201
18778468
18778936
18779749
18779865
18779912
18780043
18780060
18781815
18782018
scaffold_4, genome position

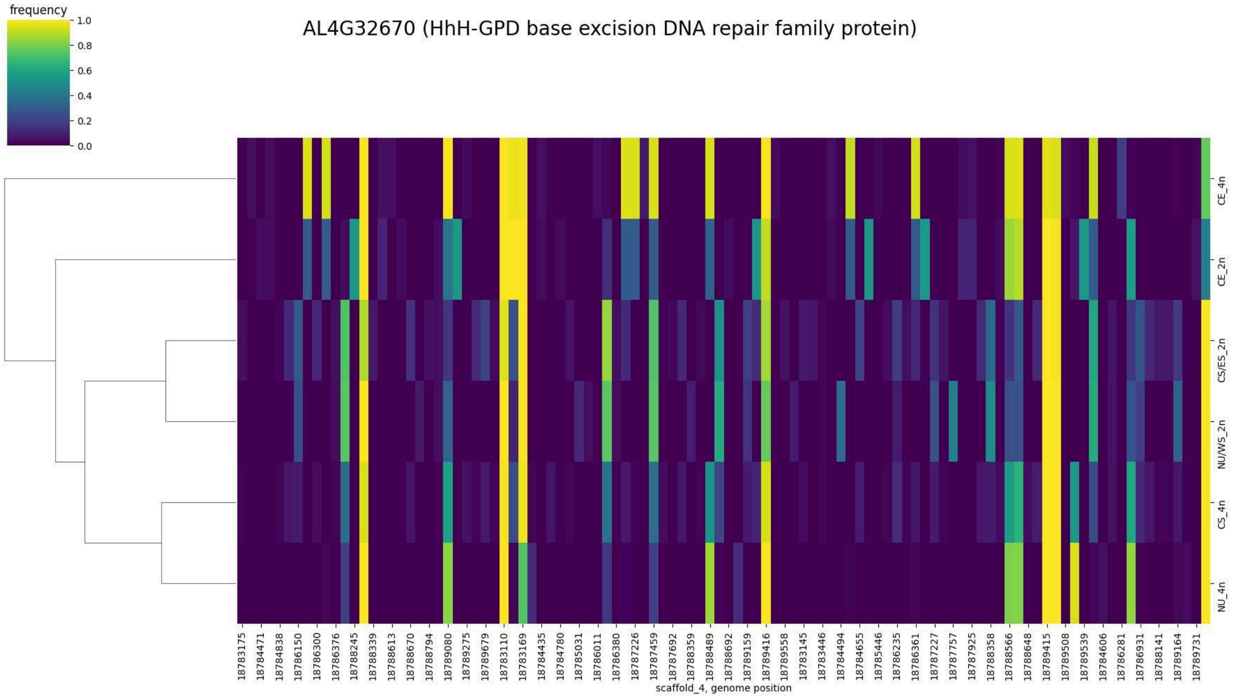
AL4G32670 (HhH-GPD base excision DNA repair family protein)
frequency
1.0
0.8
0.6
0.4
0.2
0.0
CE_4n
CE_2n
CS/ES_2n
NU/WS_2n
CS_4n
NU_4n
18783175
18784471
18784838
18786150
18786300
18786376
18788245
18788339
18788613
18788670
18788794
18789080
18789275
18789679
18783110
18783169
18784435
18784780
18785031
18786011
18786380
18787226
18787459
18787692
18788359
18788489
18788692
18789159
18789416
18789558
18783145
18783446
18784494
18784655
18785446
18786235
18786361
18787227
18787757
18787925
18788358
18788566
18788648
18789415
18789508
18789539
18784606
18786281
18786931
18788141
18789164
18789731
scaffold_4, genome position

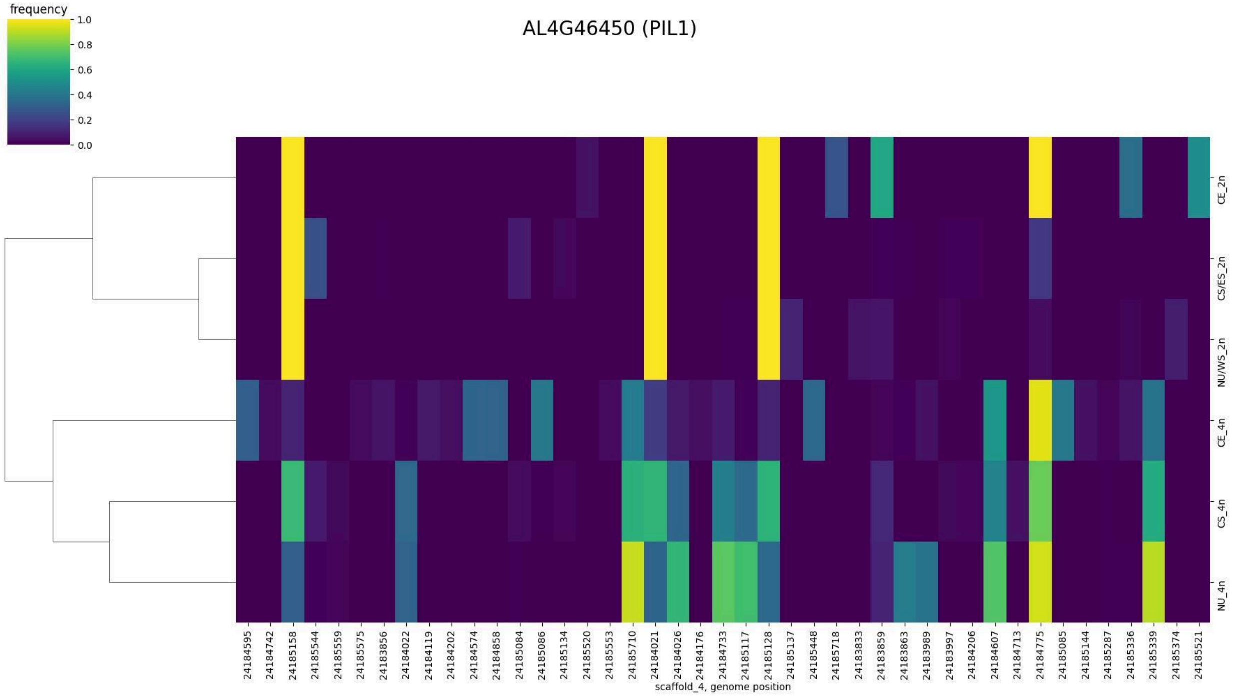
AL4G46450 (PIL1)
frequency
1.0
0.8
0.6
0.4
0.2
0.0
CE_2n
CS/ES_2n
NU/WS_2n
CE_4n
CS_4n
NU_4n
24184595
24184742
24185158
24185544
24185559
24185575
24183856
24184022
24184119
24184202
24184574
24184858
24185084
24185086
24185134
24185520
24185553
24185710
24184021
24184026
24184176
24184733
24185117
24185128
24185137
24185448
24185718
24183833
24183859
24183863
24183989
24183997
24184206
24184607
24184713
24184775
24185085
24185144
24185287
24185336
24185339
24185374
24185521
scaffold_4, genome position

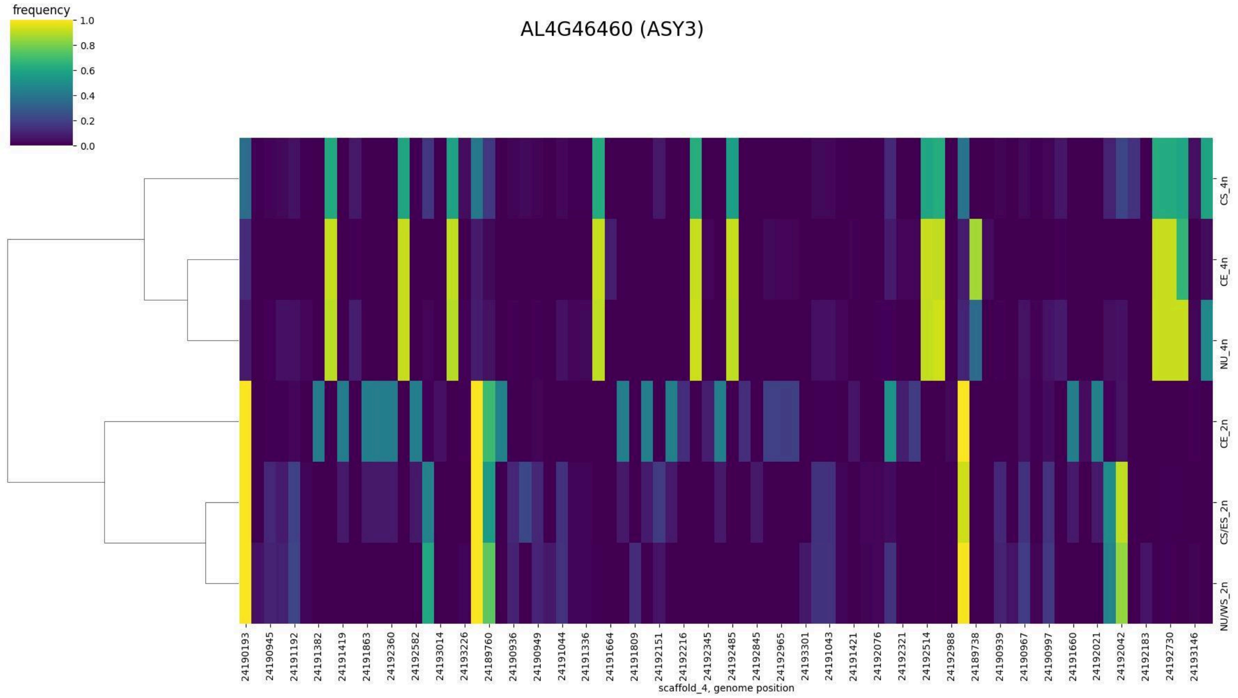
frequency
1.0
0.8
0.6
0.4
0.2
0.0
AL4G46460 (ASY3)
CS_4n
CE_4n
NU_4n
CE_2n
CS/ES_2n
NU/WS_2n
24190193
24190945
24191192
24191382
24191419
24191863
24192360
24192582
24193014
24193226
24189760
24190936
24190949
24191044
24191336
24191664
24191809
24192151
24192216
24192345
24192485
24192845
24192965
24193301
24191043
24191421
24192076
24192321
24192514
24192988
24189738
24190939
24190967
24190997
24191660
24192021
24192042
24192183
24192730
24193146
scaffold_4, genome position

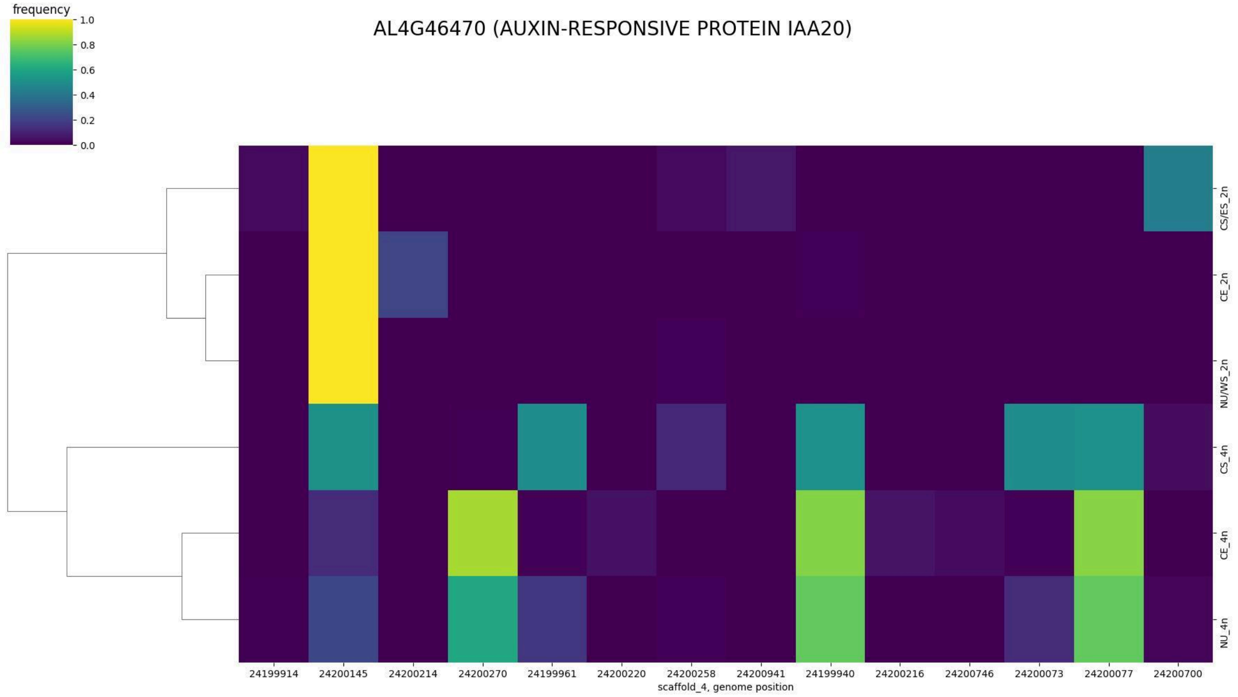
frequency
1.0
0.8
0.6
0.4
0.2
0.0
AL4G46470 (AUXIN-RESPONSIVE PROTEIN IAA20)
CS/ES_2n
CE_2n
NU/WS_2n
CS_4n
CE_4n
NU_4n
24199914
24200145
24200214
24200270
24199961
24200220
24200258
24200941
24199940
24200216
24200746
24200073
24200077
24200700
scaffold_4, genome position

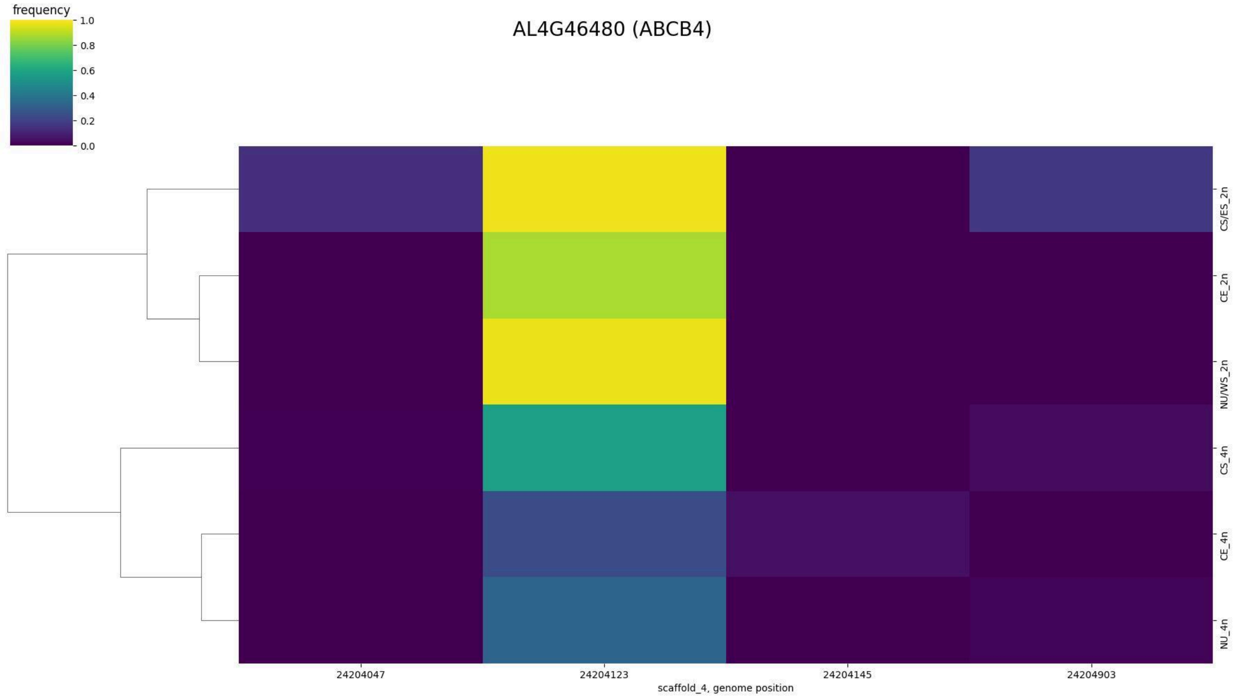
AL4G46480 (ABCB4)
frequency
1.0
0.8
0.6
0.4
0.2
0.0
CS/ES_2n
CE_2n
NU/WS_2n
CS_4n
CE_4n
NU_4n
24204047
24204123
24204145
24204903
scaffold_4, genome position

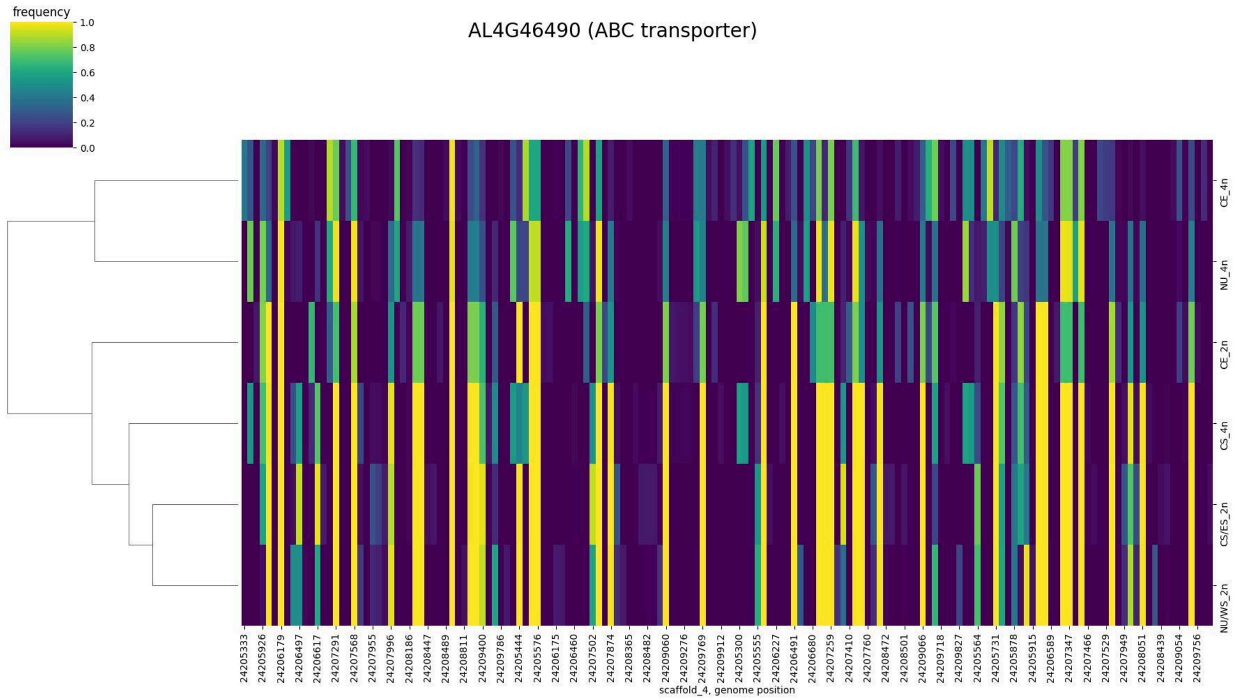
AL4G46490 (ABC transporter)
frequency
1.0
0.8
0.6
0.4
0.2
0.0
CE_4n
NU_4n
CE_2n
CS_4n
CS/ES_2n
NU/WS_2n
24205333
24205926
24206179
24206497
24206617
24207291
24207568
24207955
24207996
24208186
24208447
24208489
24208811
24209400
24209786
24205444
24205576
24206175
24206460
24207502
24207874
24208365
24208482
24209060
24209276
24209769
24209912
24205300
24205555
24206227
24206491
24206680
24207259
24207410
24207760
24208472
24208501
24209066
24209718
24209827
24205564
24205731
24205878
24205915
24206589
24207347
24207466
24207529
24207949
24208051
24208439
24209054
24209756
scaffold_4, genome position

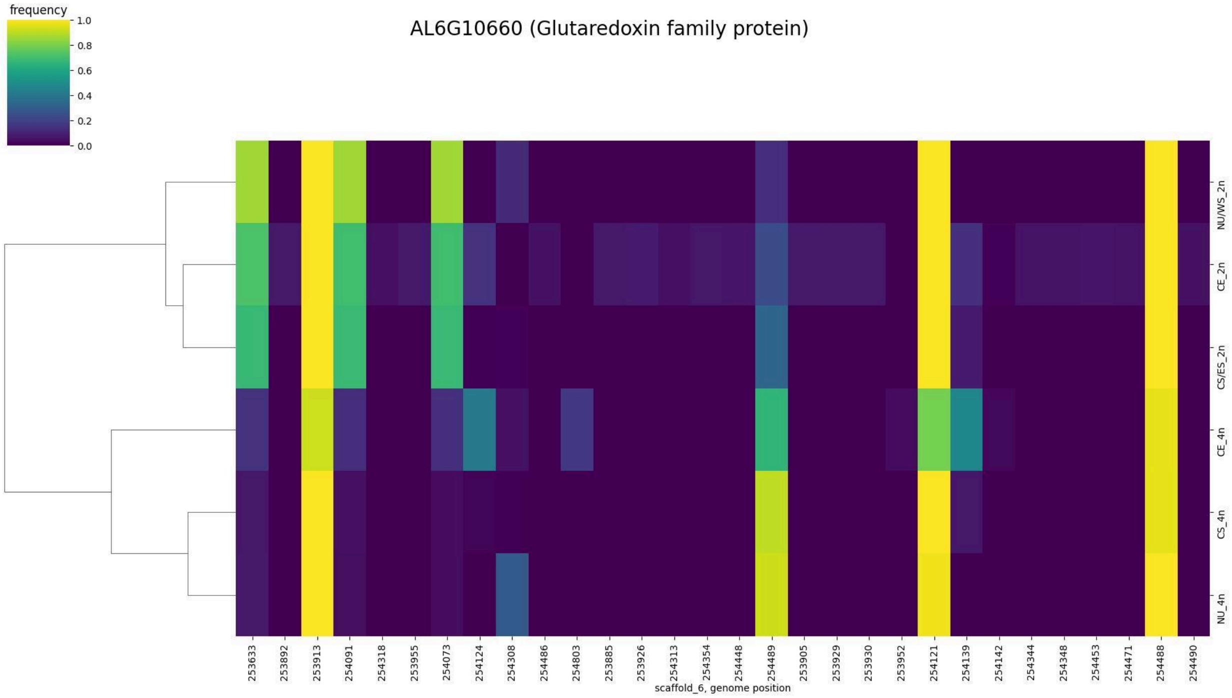
AL6G10660 (Glutaredoxin family protein)
frequency
1.0
0.8
0.6
0.4
0.2
0.0
NU/WS_2n
CE_2n
CS/ES_2n
CE_4n
CS_4n
NU_4n
253633 253892 253913 254091 254318 253955 254073 254124 254308 254486 254803 253885 253926 254313 254354 254448 254489 253905 253929 253930 253952 254121 254139 254142 254344 254348 254453 254471 254488 254490
scaffold_6, genome position

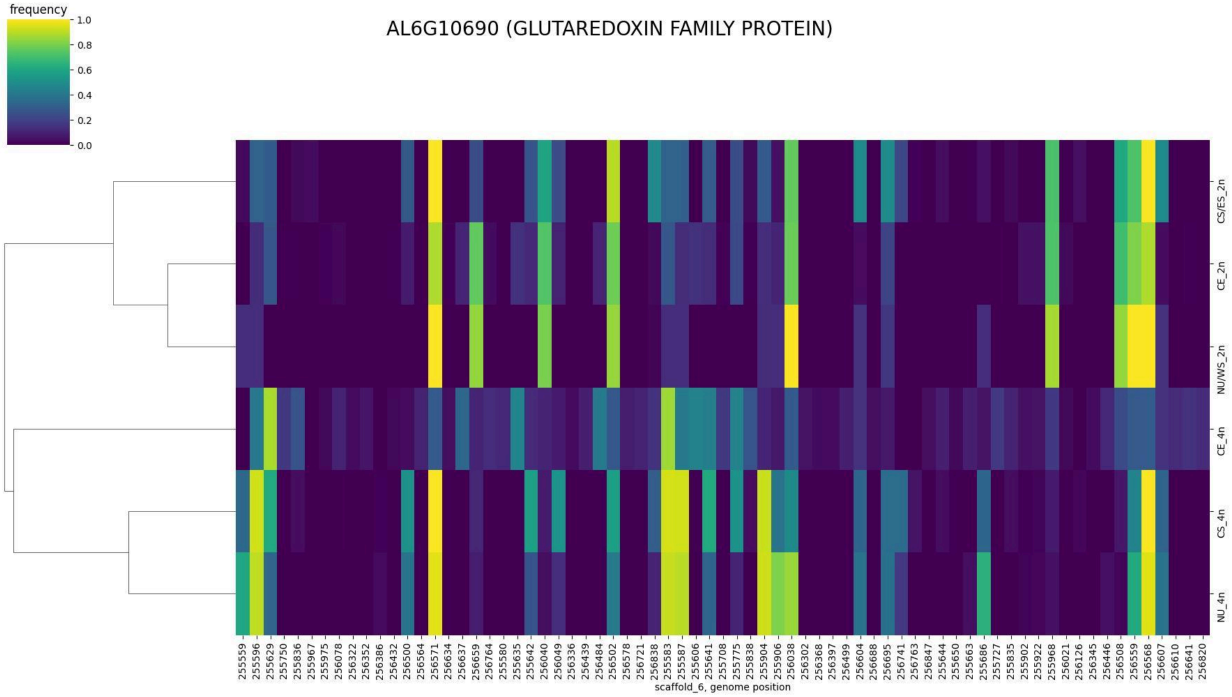
AL6G10690 (GLUTAREDOXIN FAMILY PROTEIN)
frequency
1.0
0.8
0.6
0.4
0.2
0.0
CS/ES_2n
CE_2n
NU/WS_2n
CE_4n
CS_4n
NU_4n
scaffold_6, genome position

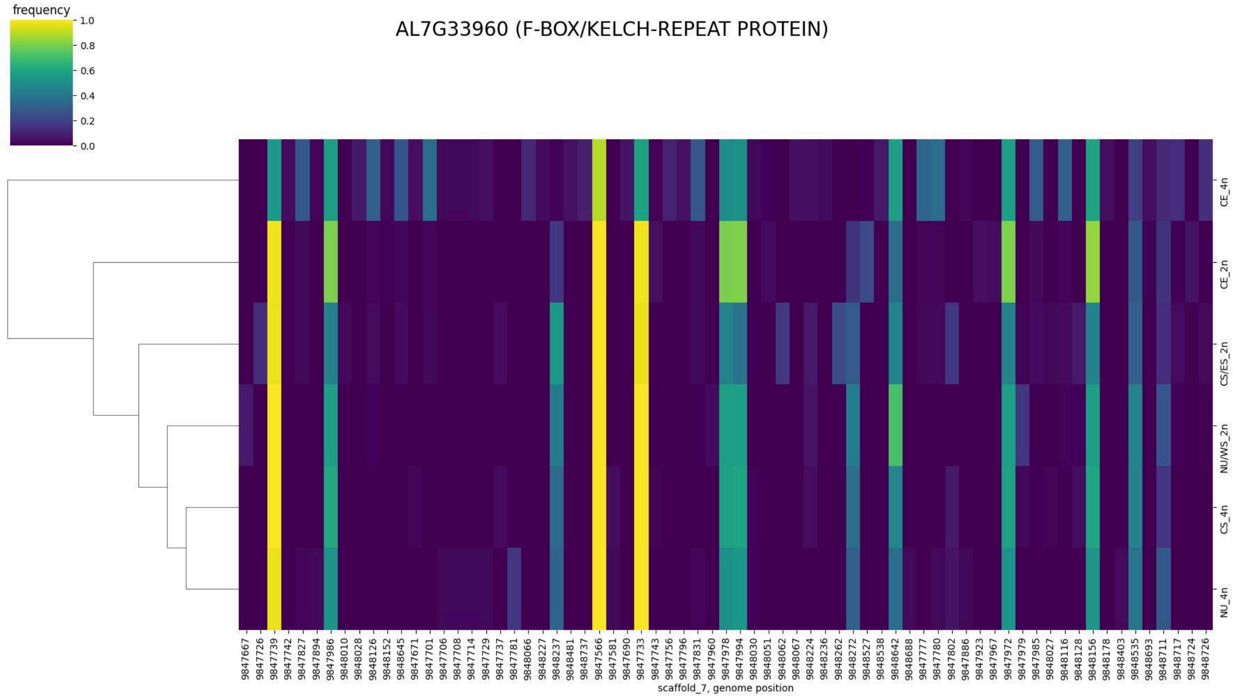
AL7G33960 (F-BOX/KELCH-REPEAT PROTEIN)
frequency
1.0
0.8
0.6
0.4
0.2
0.0
CE_4n
CE_2n
CS/ES_2n
NU/WS_2n
CS_4n
NU_4n
9847667 9847726 9847739 9847742 9847827 9847894 9847986 9848010 9848028 9848126 9848152 9848645 9847671 9847701 9847706 9847708 9847714 9847729 9847737 9847781 9848066 9848227 9848237 9848481 9848737 9847566 9847581 9847690 9847733 9847743 9847756 9847796 9847831 9847960 9847978 9847994 9848030 9848051 9848062 9848067 9848224 9848236 9848262 9848272 9848527 9848538 9848642 9848688 9847777 9847780 9847802 9847886 9847923 9847967 9847972 9847979 9847985 9848027 9848116 9848128 9848156 9848178 9848403 9848535 9848693 9848711 9848717 9848724 9848726
scaffold_7, genome position

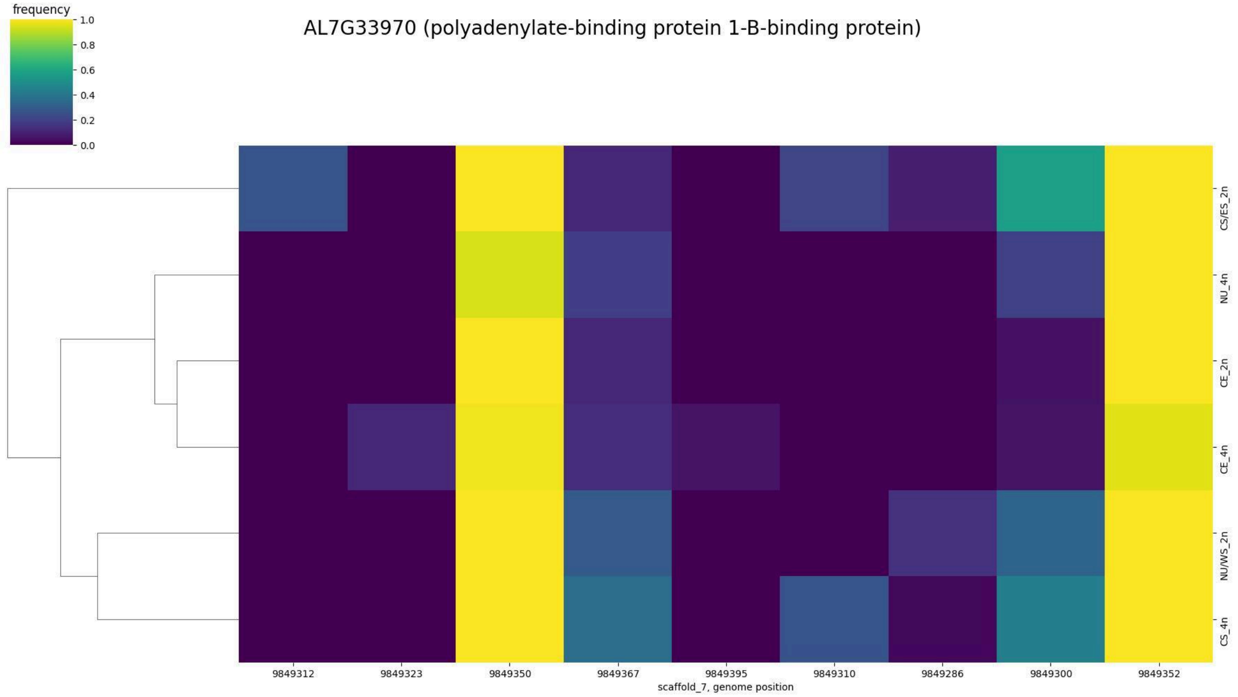
AL7G33970 (polyadenylate-binding protein 1-B-binding protein)
frequency
1.0
0.8
0.6
0.4
0.2
0.0
CS/ES_2n
NU_4n
CE_2n
CE_4n
NU/WS_2n
CS_4n
9849312 9849323 9849350 9849367 9849395 9849310 9849286 9849300 9849352
scaffold_7, genome position

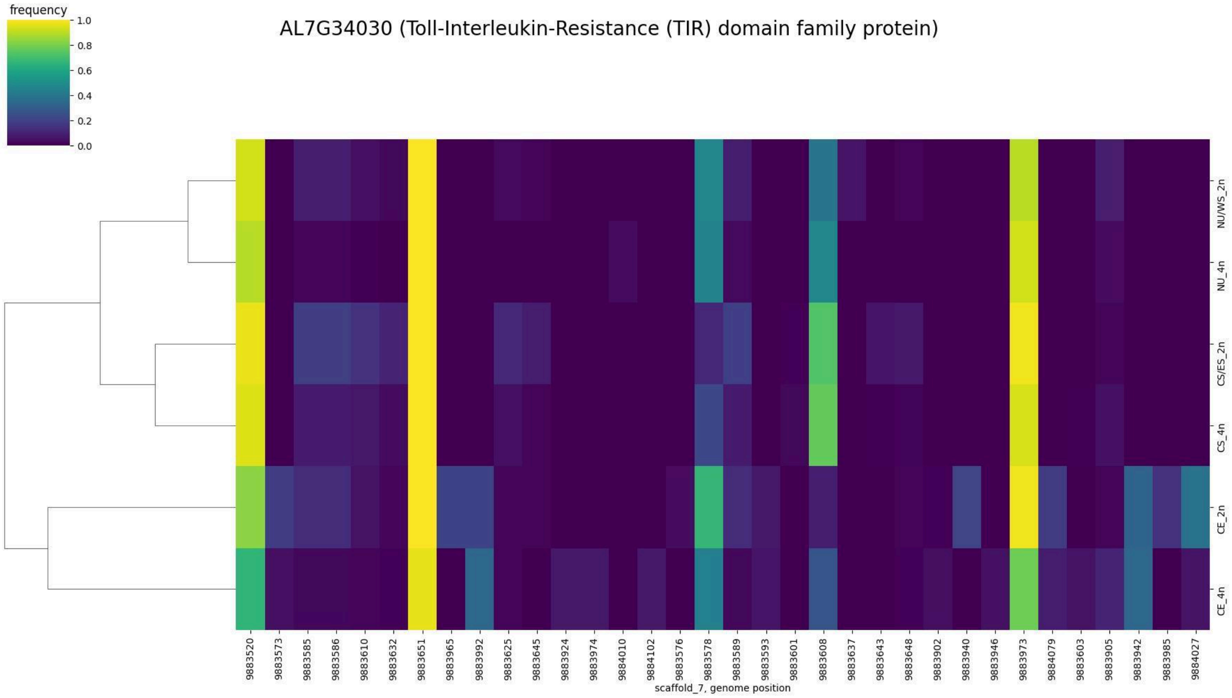

frequency
1.0
0.8
0.6
0.4
0.2
0.0
AL7G34030 (Toll-Interleukin-Resistance (TIR) domain family protein)
NU/WS_2n
NU_4n
CS/ES_2n
CS_4n
CE_2n
CE_4n
9883520
9883573
9883585
9883586
9883610
9883632
9883651
9883965
9883992
9883625
9883645
9883924
9883974
9884010
9884102
9883576
9883578
9883589
9883593
9883601
9883608
9883637
9883643
9883648
9883902
9883940
9883946
9883973
9884079
9883603
9883905
9883942
9883985
9884027
scaffold_7, genome position

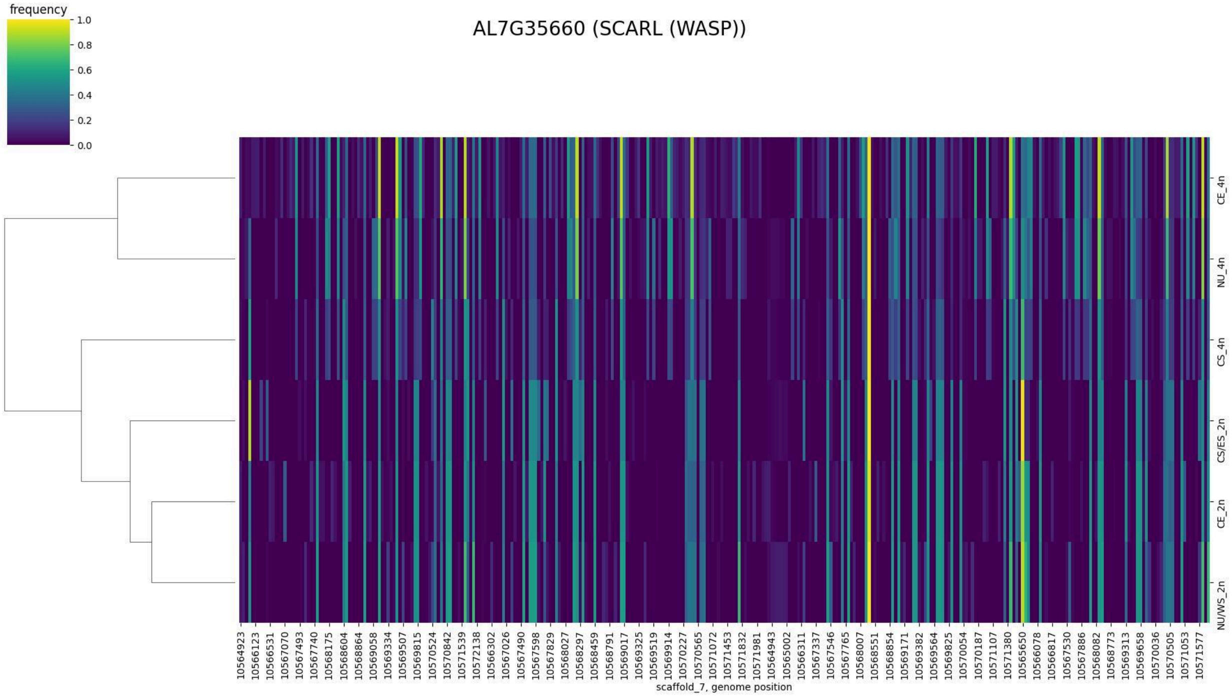

frequency
1.0
0.8
0.6
0.4
0.2
0.0
AL7G35660 (SCARL (WASP))
CE_4n
NU_4n
CS_4n
CS/ES_2n
CE_2n
NU/WS_2n
scaffold_7, genome position

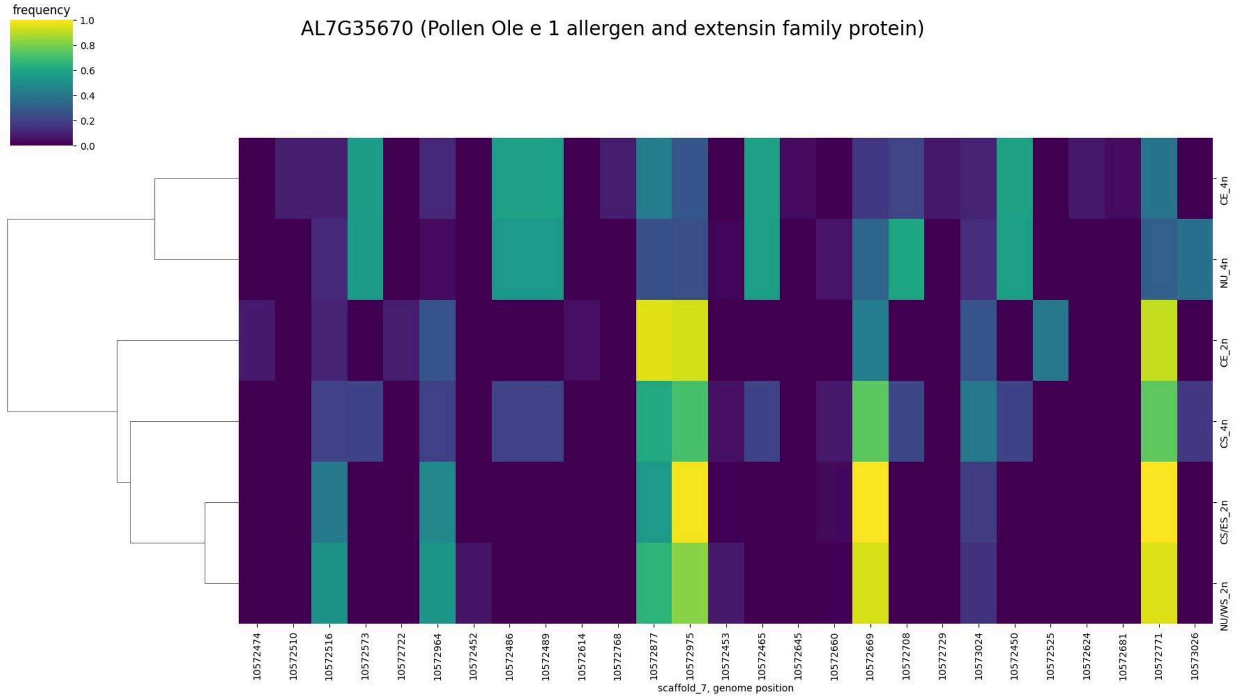

AL7G35670 (Pollen Ole e 1 allergen and extensin family protein)
frequency
1.0
0.8
0.6
0.4
0.2
0.0
CE_4n
NU_4n
CE_2n
CS_4n
CS/ES_2n
NU/WS_2n
10572474
10572510
10572516
10572573
10572722
10572964
10572452
10572486
10572489
10572614
10572768
10572877
10572975
10572453
10572465
10572645
10572660
10572669
10572708
10572729
10573024
10572450
10572525
10572624
10572681
10572771
10573026
scaffold_7, genome position

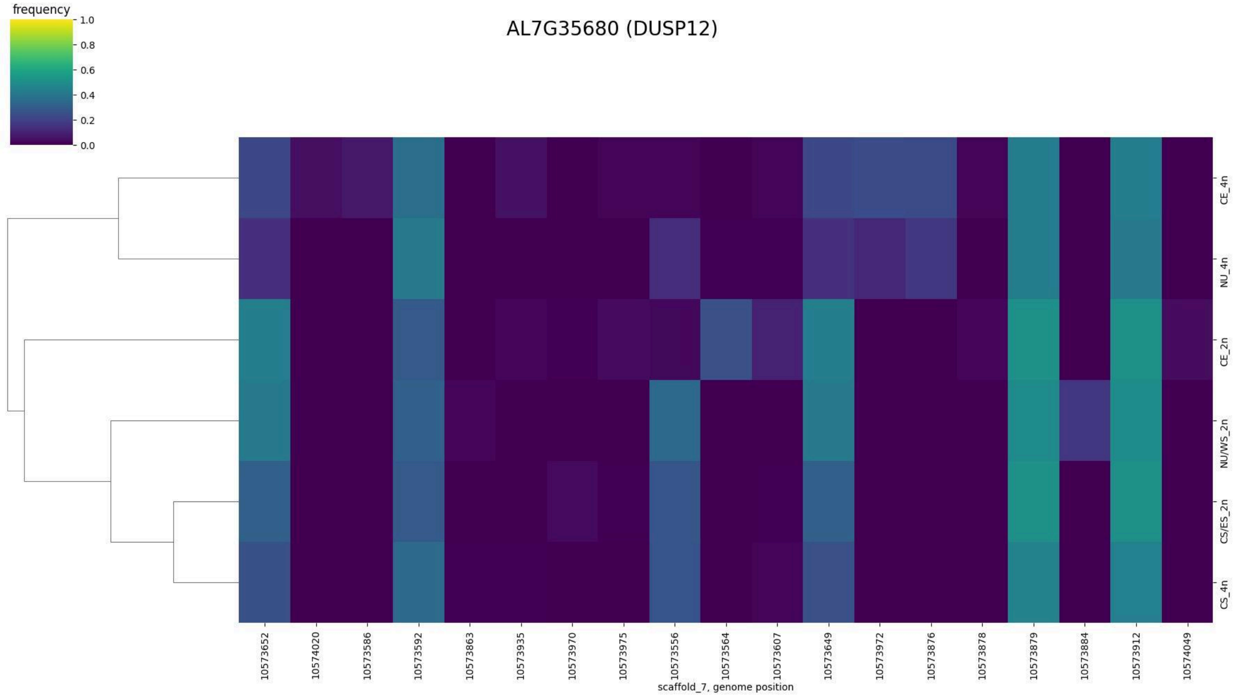

AL7G35680 (DUSP12)
frequency
1.0
0.8
0.6
0.4
0.2
0.0
CE_4n
NU_4n
CE_2n
NU/WS_2n
CS/ES_2n
CS_4n
10573652
10574020
10573586
10573592
10573863
10573935
10573970
10573975
10573556
10573564
10573607
10573649
10573972
10573876
10573878
10573879
10573884
10573912
10574049
scaffold_7, genome position

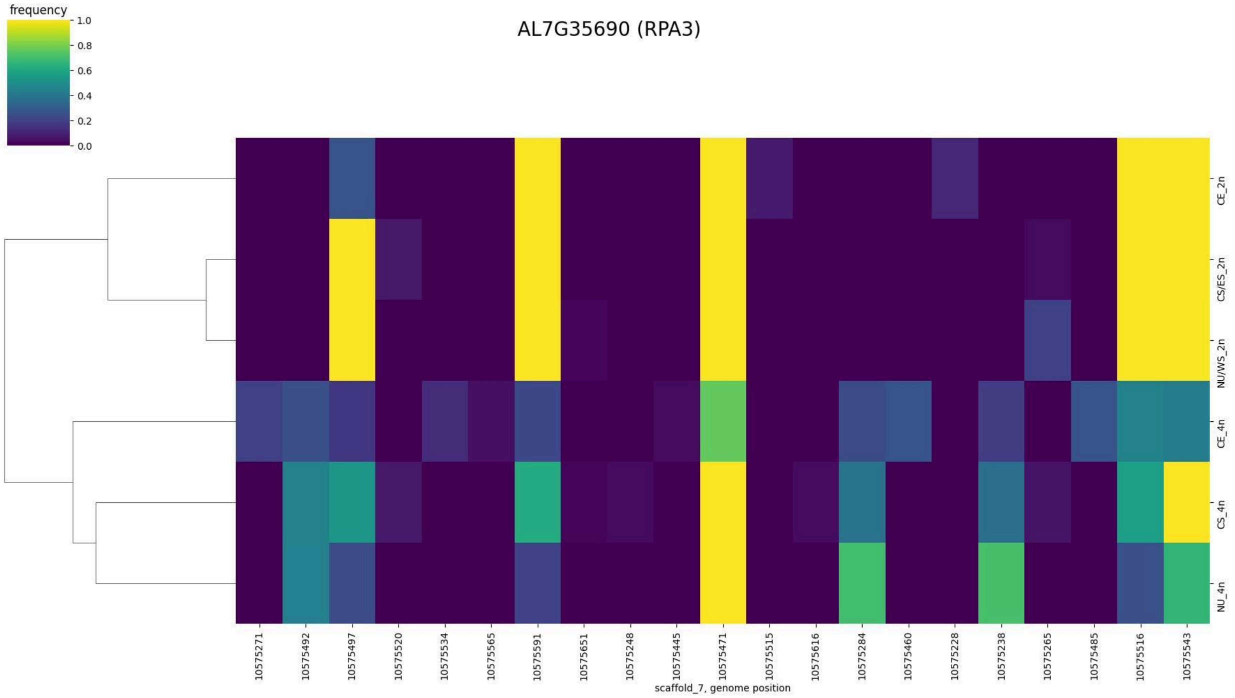
frequency
1.0
0.8
0.6
0.4
0.2
0.0
AL7G35690 (RPA3)
CE_2n
CS/ES_2n
NU/WS_2n
CE_4n
CS_4n
NU_4n
10575271
10575492
10575497
10575520
10575534
10575565
10575591
10575651
10575248
10575445
10575471
10575515
10575616
10575284
10575460
10575228
10575238
10575265
10575485
10575516
10575543
scaffold_7, genome position

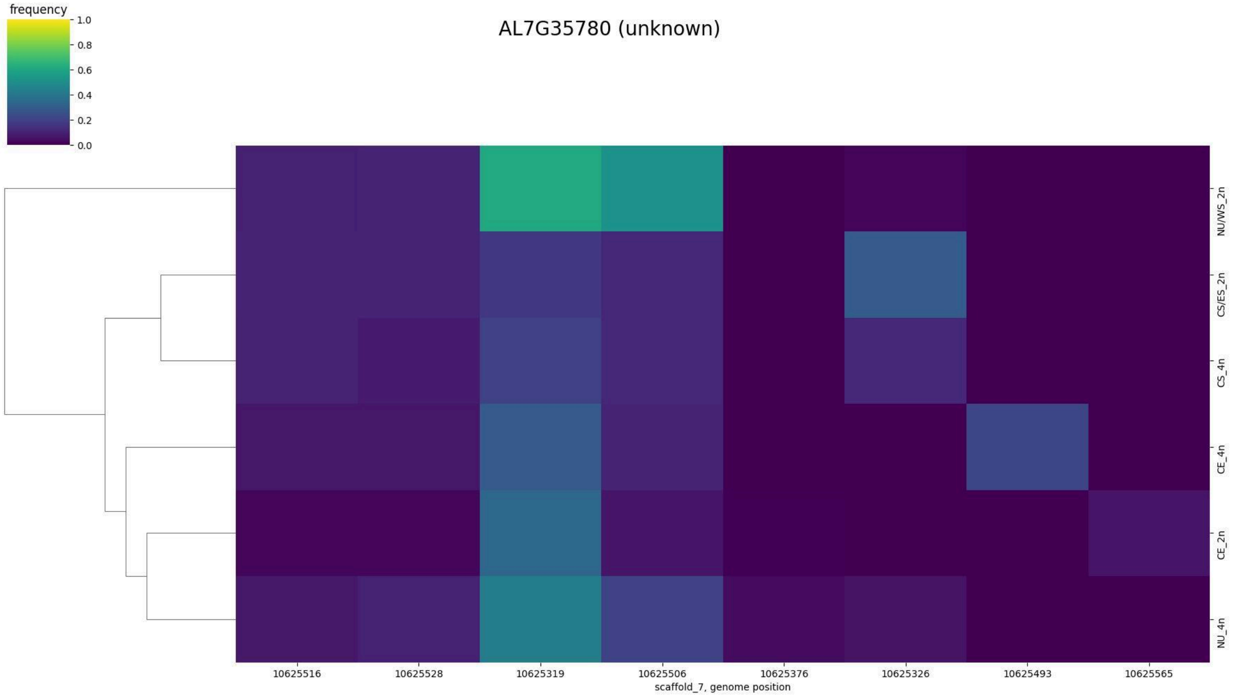
frequency
1.0
0.8
0.6
0.4
0.2
0.0
AL7G35780 (unknown)
NU/WS_2n
CS/ES_2n
CS_4n
CE_4n
CE_2n
NU_4n
10625516
10625528
10625319
10625506
10625376
10625326
10625493
10625565
scaffold_7, genome position

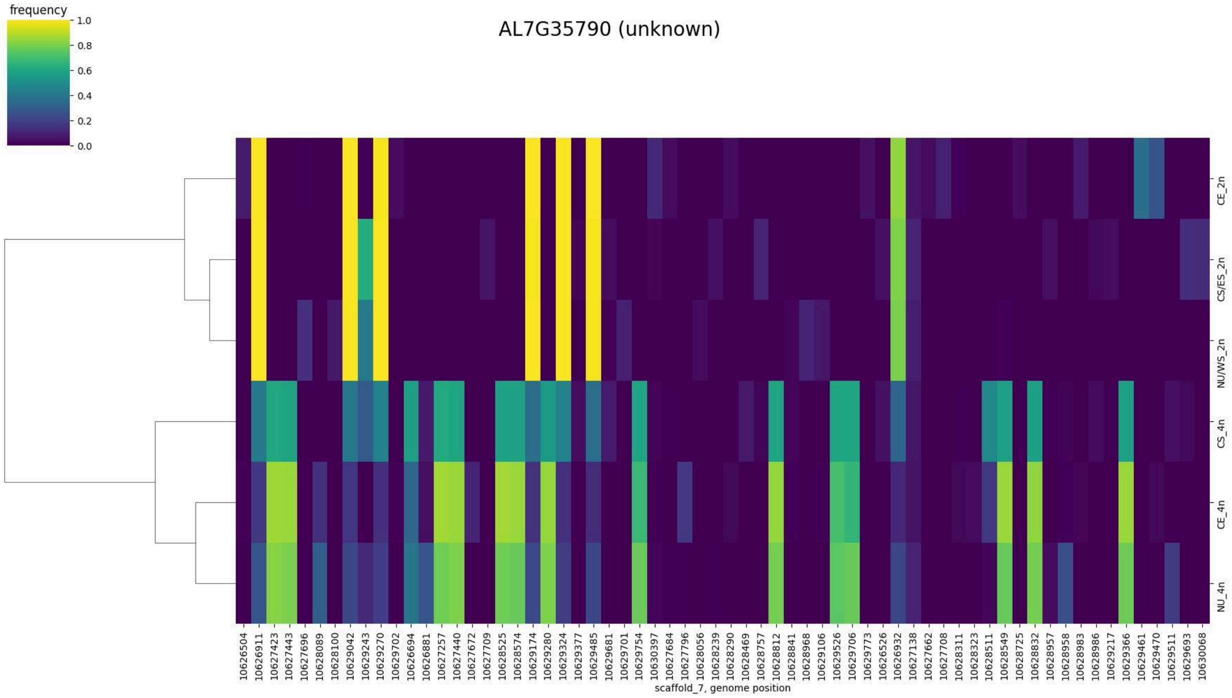
AL7G35790 (unknown)
frequency
1.0
0.8
0.6
0.4
0.2
0.0
CE_2n
CS/ES_2n
NU/WS_2n
CS_4n
CE_4n
NU_4n
scaffold_7, genome position

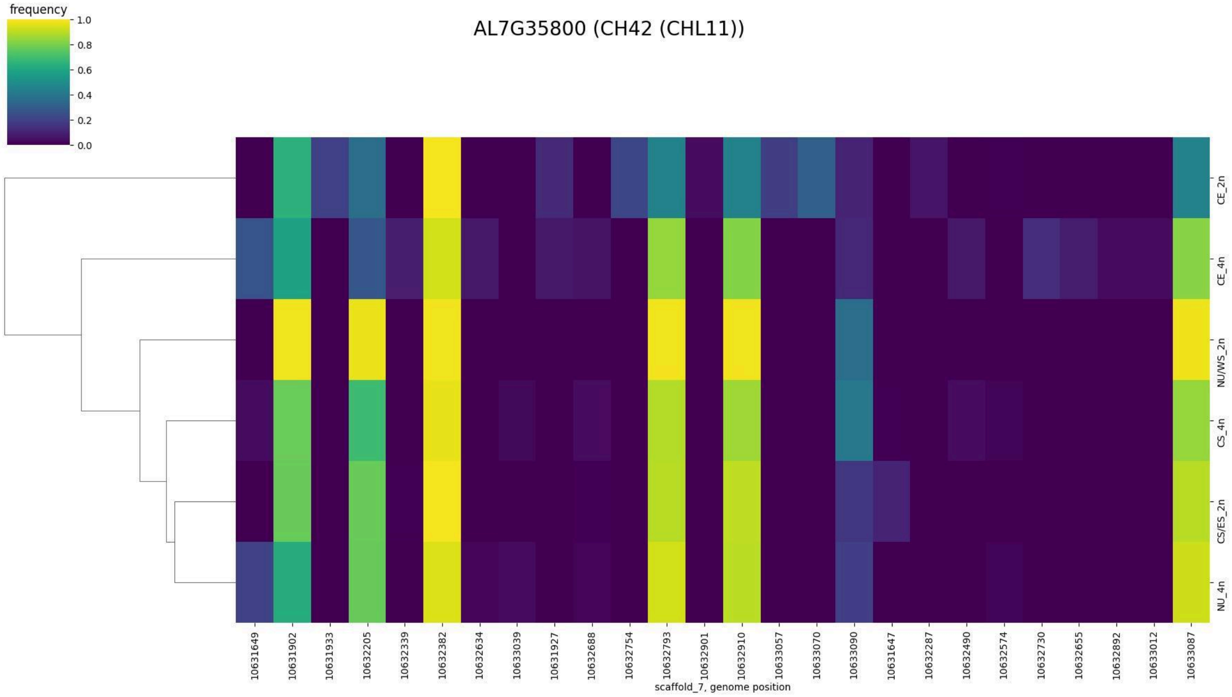
AL7G35800 (CH42 (CHL11))
frequency
1.0
0.8
0.6
0.4
0.2
0.0
CE_2n
CE_4n
NU/WS_2n
CS_4n
CS/ES_2n
NU_4n
scaffold_7, genome position

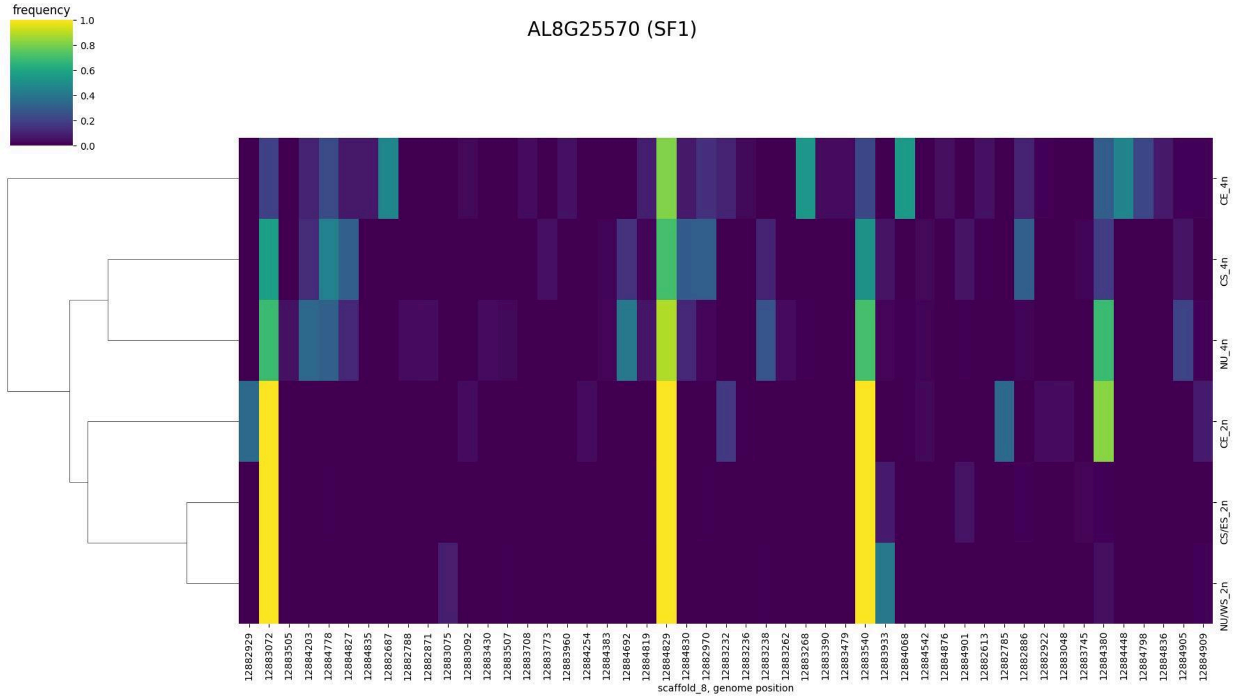
frequency
1.0
0.8
0.6
0.4
0.2
0.0
AL8G25570 (SF1)
CE_4n
CS_4n
NU_4n
CE_2n
CS/ES_2n
NU/WS_2n
scaffold_8, genome position

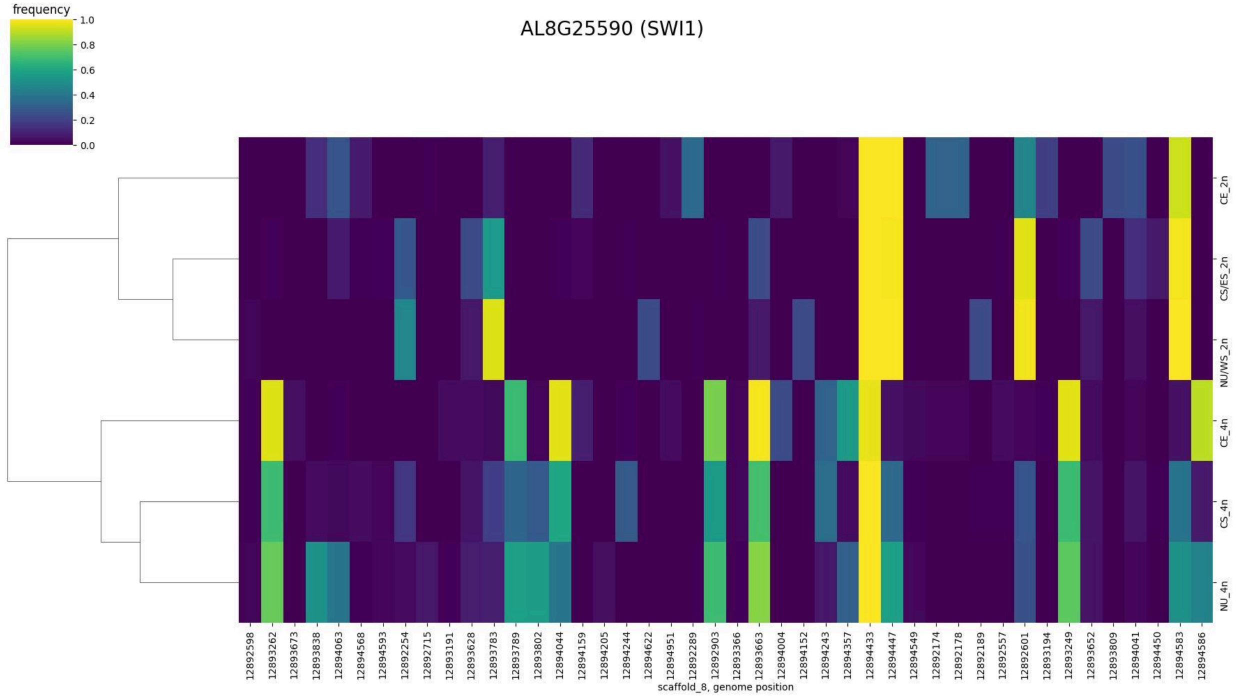
frequency
1.0
0.8
0.6
0.4
0.2
0.0
AL8G25590 (SWI1)
CE_2n
CS/ES_2n
NU/WS_2n
CE_4n
CS_4n
NU_4n
scaffold_8, genome position

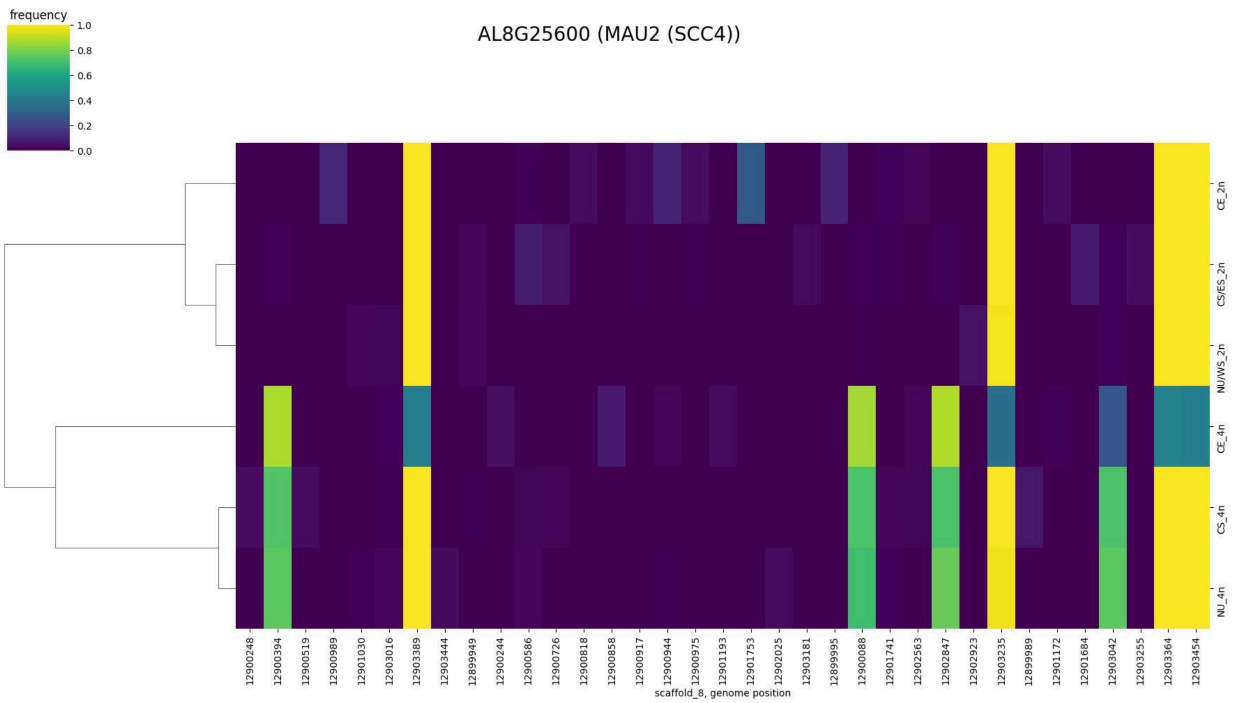
AL8G25600 (MAU2 (SCC4))
frequency
1.0
0.8
0.6
0.4
0.2
0.0
CE_2n
CS/ES_2n
NU/WS_2n
CE_4n
CS_4n
NU_4n
12900248
12900394
12900519
12900989
12901030
12903016
12903389
12903444
12899949
12900244
12900586
12900726
12900818
12900858
12900917
12900944
12900975
12901193
12901753
12902025
12903181
12899995
12900088
12901741
12902563
12902847
12902923
12903235
12899989
12901172
12901684
12903042
12903255
12903364
12903454
scaffold_8, genome position

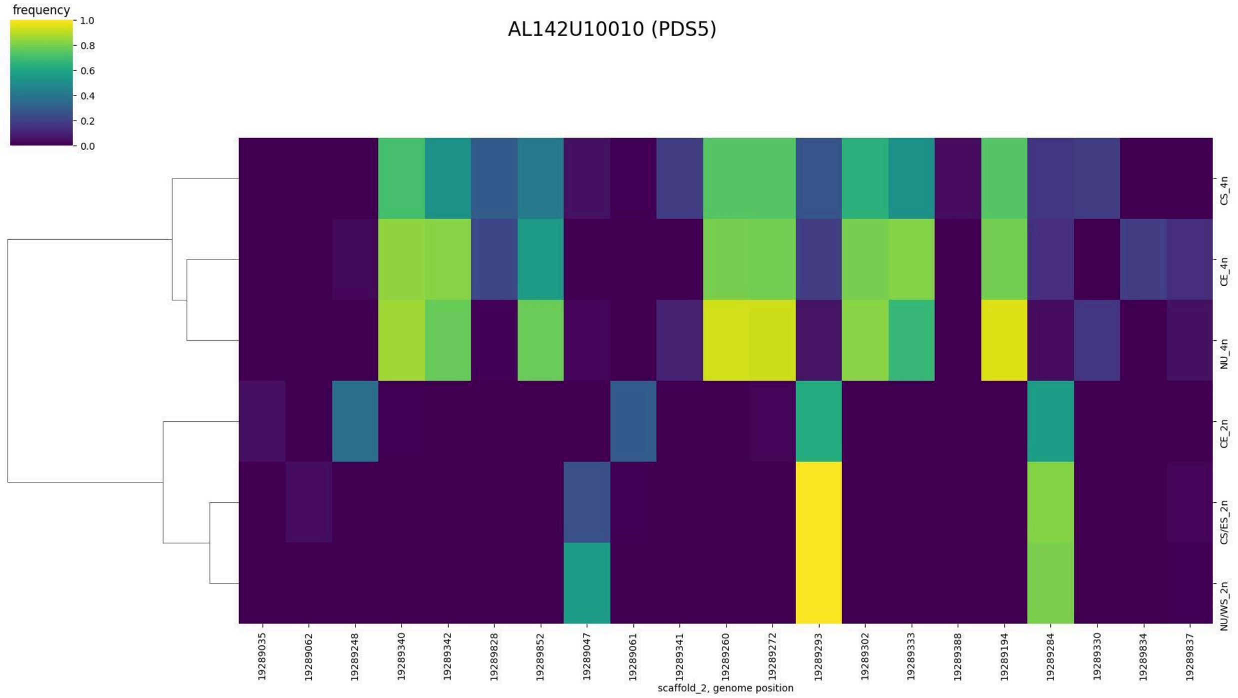

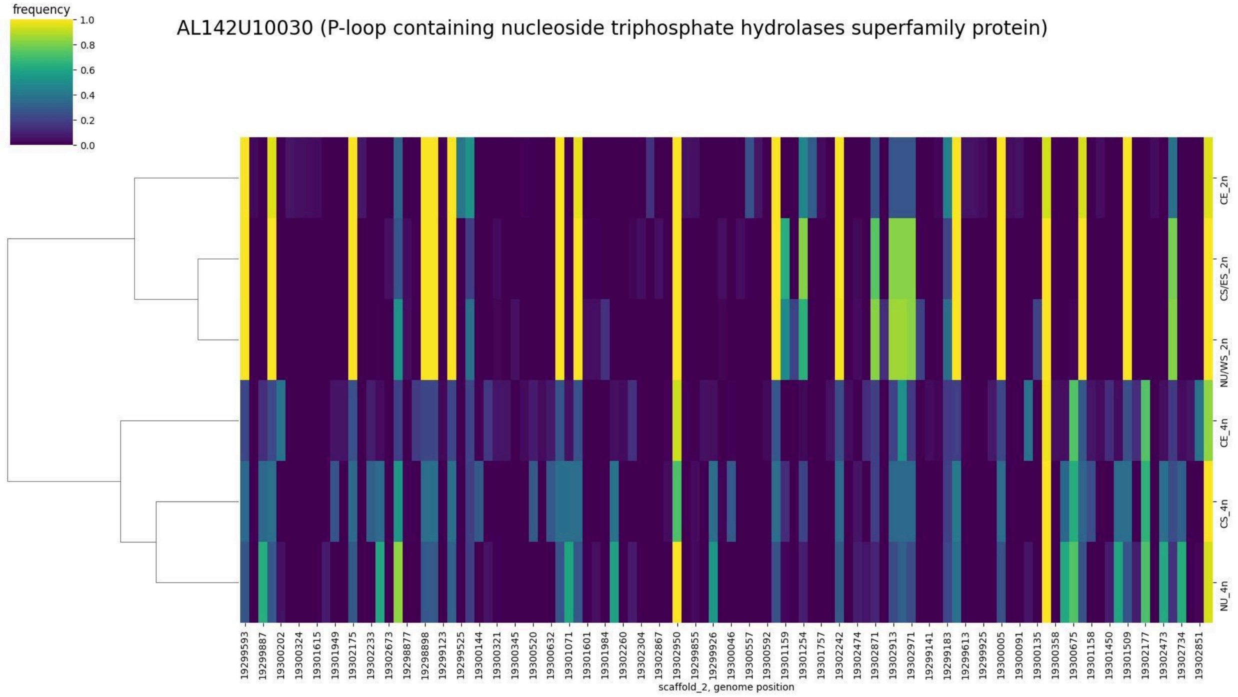

Supplementary Figure 7. Biallelic SNP frequency heatmap for each introgressed gene. Hierarchical clustering based on distance shown on the left side of the heatmap.

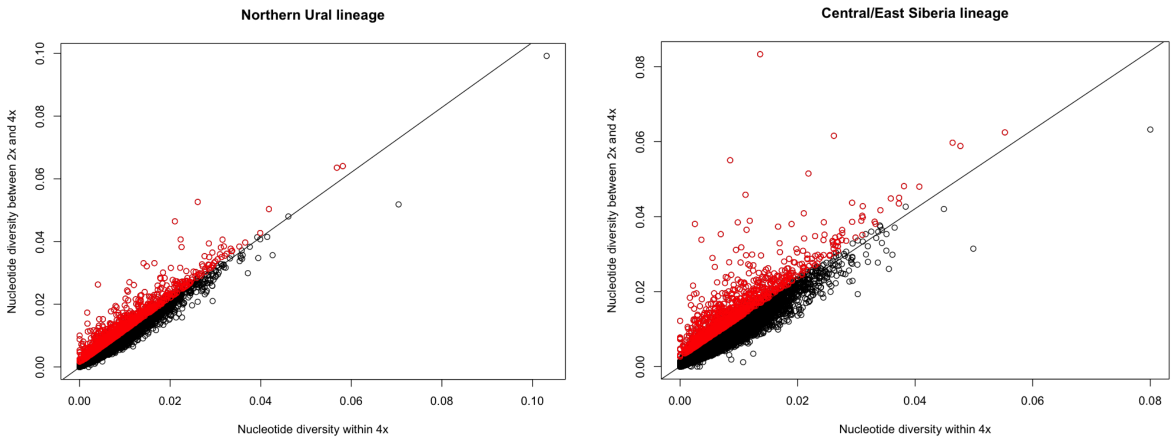

Supplementary Figure 7. Nucleotide diversity within tetraploids (π) and between tetraploids and diploids (dxy). Nucleotide diversity is calculated per-gene, so that each dot represents a single gene. Outlier genes colored red for each comparison.

Supplementary Table 1 - Per-population calculations of pi, per-chromosome and averaged across the genome.

| Pop | n | Ploidy | Scaff 1 | Scaff 2 | Scaff 3 | Scaff 4 | Scaff 5 | Scaff 6 | Scaff 7 | Scaff 8 | Genome average |
|---|---|---|---|---|---|---|---|---|---|---|---|
| WS | 7 | 4 | 0.0046813 | 0.0058471 | 0.0044556 | 0.0054778 | 0.0050703 | 0.0042065 | 0.0049327 | 0.0049792 | 0.0049563537 |
| PU_6 | 8 | 4 | 0.0052483 | 0.0059976 | 0.0046710 | 0.005990 | 0.0059692 | 0.0047288 | 0.0055707 | 0.005536 | 0.00546408 |
| NT_16 | 5 | 2 | 0.0042319 | 0.0061000 | 0.0041937 | 0.0055608 | 0.0050671 | 0.004080 | 0.0049527 | 0.0051636 | 0.0049188462 |
| TE_10 | 9 | 2 | 0.0039169 | 0.005482 | 0.0041352 | 0.0054400 | 0.0053192 | 0.0037957 | 0.0047216 | 0.0047287 | 0.004692492 |
| TE_11 | 10 | 2 | 0.0039079 | 0.0053864 | 0.004015 | 0.005095 | 0.0049126 | 0.0037345 | 0.0045031 | 0.0046341 | 0.00452371 |
| TE_3 | 9 | 2 | 0.0040873 | 0.0057677 | 0.0038921 | 0.0049703 | 0.0048731 | 0.003762 | 0.0047304 | 0.0045185 | 0.00457525 |
| TE_4 | 10 | 2 | 0.0040584 | 0.0059398 | 0.0040612 | 0.0050558 | 0.0047924 | 0.0038124 | 0.0047723 | 0.0045181 | 0.004626352 |
| TE_5 | 10 | 2 | 0.004034 | 0.0056828 | 0.0037725 | 0.0049740 | 0.0047831 | 0.0037380 | 0.0046142 | 0.0046314 | 0.0045288412 |
| TE_7 | 9 | 2 | 0.0042261 | 0.0057393 | 0.0038766 | 0.0052549 | 0.0051601 | 0.0036793 | 0.0045437 | 0.0044164 | 0.004612 |
| TE_8 | 10 | 2 | 0.0038363 | 0.0049451 | 0.0033700 | 0.0040193 | 0.0046799 | 0.0032951 | 0.004185 | 0.004250 | 0.0040727137 |

Supplementary Table 2 - Model comparison and parameter estimates from demographic modeling. Population sizes in no. of alleles, divergence time in generations.

| **Central Siberian diploids & tetraploids** | **Input alleles (16)** | **Input alleles (32)** | | | | | | | |
|---|---|---|---|---|---|---|---|---|---|
| | NPOP1 | NPOP2 | NANC | TDIV | MIG1 | Max EstLhood | Max ObsLhood | AIC | Parameters |
| Simple divergence | 41752 | 9655 | 16468 | 2845 | - | -1,131,335 | -1,098,814 | 5,209,998.21 | 4 |
| **Divergence + gene flow (dip -> tet)** | **38373** | **5814** | **14438** | **5973** | **1.44E-04** | **-1,123,698** | **-1,098,814** | **5,174,832.53** | **5** |
| 2.5% | 36582.28 | 5473.604 | 13973.06 | 5657.09 | 1.49E-04 | | | | |
| 97.5% | 36986.16 | 5538.786 | 14086.44 | 5729.40 | 1.51E-04 | | | | |
| mean value CI | 36784.22 | 5506.195 | 14029.75 | 5693.25 | 1.50E-04 | | | | |
| **Northern Ural diploids & tetraploids** | **Input alleles (10)** | **Input alleles (20)** | | | | | | | |
| | NPOP1 | NPOP2 | NANC | TDIV | MIG1 | Max EstLhood | Max ObsLhood | AIC | Parameters |
| Simple divergence | 34008 | 14176 | 16159 | 5355 | - | -1,055,296 | -1,039,996 | 4,859,826.60 | 4 |
| **Divergence + gene flow (dip -> tet)** | **29636** | **10859** | **14272** | **8660** | **5.44E-05** | **-1,052,262** | **-1,039,996** | **4,845,856.51** | **5** |
| 2.5% | 30185.72 | 10332.68 | 13573.33 | 8080.308 | 5.37E-05 | | | | |
| 97.5% | 30462.13 | 10436.62 | 13678.58 | 8150.242 | 5.45E-05 | | | | |
| mean value CI | 30323.92 | 10384.65 | 13625.95 | 8115.275 | 5.41E-05 | | | | |

Supplementary Table 3 - Published and assembled S-alleles used in this study.

| S-allele | Corresponding name | Accession | Reference |
|---|---|---|---|
| H2018 | CgrSRK45 | MT592980.1 | Neuffer et al., 2023 |
| H3009 | AhSRK30 | EU878015.1 | Castric et al., 2008 |
| H3012 | CgrSRK11 | MT592933.1 | Neuffer et al., 2023 |
| H3014 | CgrSRK19 | MT592944.1 | Neuffer et al., 2023 |
| H3021 | - | - | supplementary data 3 |
| H3024 | - | - | supplementary data 3 |

| | | | |
|---|---|---|---|
| H4021 | CgrSRK56 | MT592996.1 | Neuffer et al., 2023 |
| H4031 | CgrSRK28 | MT592956.1 | Neuffer et al., 2023 |
| H4042 | - | - | supplementary data 3 |
| H4043 | - | - | supplementary data 3 |
